# Countering Cross-Individual Variance in Event Related Potentials with Functional Profiling

**DOI:** 10.1101/455030

**Authors:** Abdulmajeed Alsufyani, Alexia Zoumpoulaki, Marco Filetti, Dirk P. Janssen, Howard Bowman

## Abstract

In this study, a simple method called the Weight Template (WT) is proposed for classifying brain responses of individuals into deceiving and non-deceiving. The performance of our method was evaluated on a number of identity deception data sets and on artificial EEG data sets. A comparison was made with a standard method used to measure P3 presence, called Peak-to-Peak. In the real experimental data, the WT showed higher performance in terms of sensitivity and specificity. In the artificial EEG data, in ERPs with low Signal-to-Noise Ratio (SNR), the WT was more resistant to noise and provided more accurate measures.

**Highlights:**

- We propose a method for classifying ERPs of the P3 Concealed Information Tests.
- The new method is based on computing a weighted template (WT) for each individual.
- The WT is used to characterize the shape of each individual’s P3.
- The performance of the WT was compared with the standard Peak-to-Peak method.
- The WT showed higher performance in terms of sensitivity and specificity.

## 1 Introduction

In Event Related Potential (ERP) research, per-individual inference (e.g., when detecting that a single person is lying) and standard cross-individual group-level inference are made harder by the presence of considerable cross-individual variance in respect of the latency and form of ERP components. In particular, an individual’s ERP pattern may be very different to the mean population pattern, as approximated by grand averages. However, most ERP research is strongly wedded to a notion of fixed-form components, e.g., that the P3b is a positive deflection in a window from 250 to 600 ms after stimulus presentation. This fixed-form is often clearly evident at the grand average-level, but much less so for some individuals’ ERPs.

The emphasis on fixed-form components has been a central principle of ERP research, which has led to many clear and compelling cognitive interpretations, e.g., “working memory update occurs between 300 and 500 ms after stimulus presentation and following sensory processing (PI-NI complex), etc.” However, in the presence of individual differences in ERP-form, it may sacrifice considerable statistical power.

The fixed-form components approach also reflects a wholely justified concern not to fish and thereby to prevent increase in the effective number of comparisons. The problem is that, under classical statistical inference, *tailoring* identification-parameters, such as, window placement, to the deflections in the observed ERP contrast is post hoc fishing. That is, even under the null hypothesis, i.e., when an ERP contrast really does not contain any hypothesis-distinguishing difference, there will be window placements that will isolate statistically significant differences. Moreover, such differences will be identifiable under tailoring with a probability higher than the alpha level, potentially inflating the type one error rate considerably; see for example (Kilner, 2013).

Reliance on fixed-form components resolves this issue if that form, and thus window-placement, is justified a priori, e.g., by some previous published study. This then is a statistically safe approach, in which the type one error rate is controlled. However, variability across individuals is likely to ensure a loss of statistical power, i.e., an inflation of the type two error rate. As an illustration, an individual in which the P3 is accelerated might contribute zero (or even a negative number) in a condition of a cross-participant test, because the negativity that often follows the P3 positivity also fell in the a priori fixed window (see Figure 1). However, in such situations, “real” effects may be present, even though they cannot be found under a fixed-form components assumption; and our example demonstrates this: the individual with an accelerated P3 does seem to be exhibiting a P3, it is just atypical in form.

**Figure 1.**
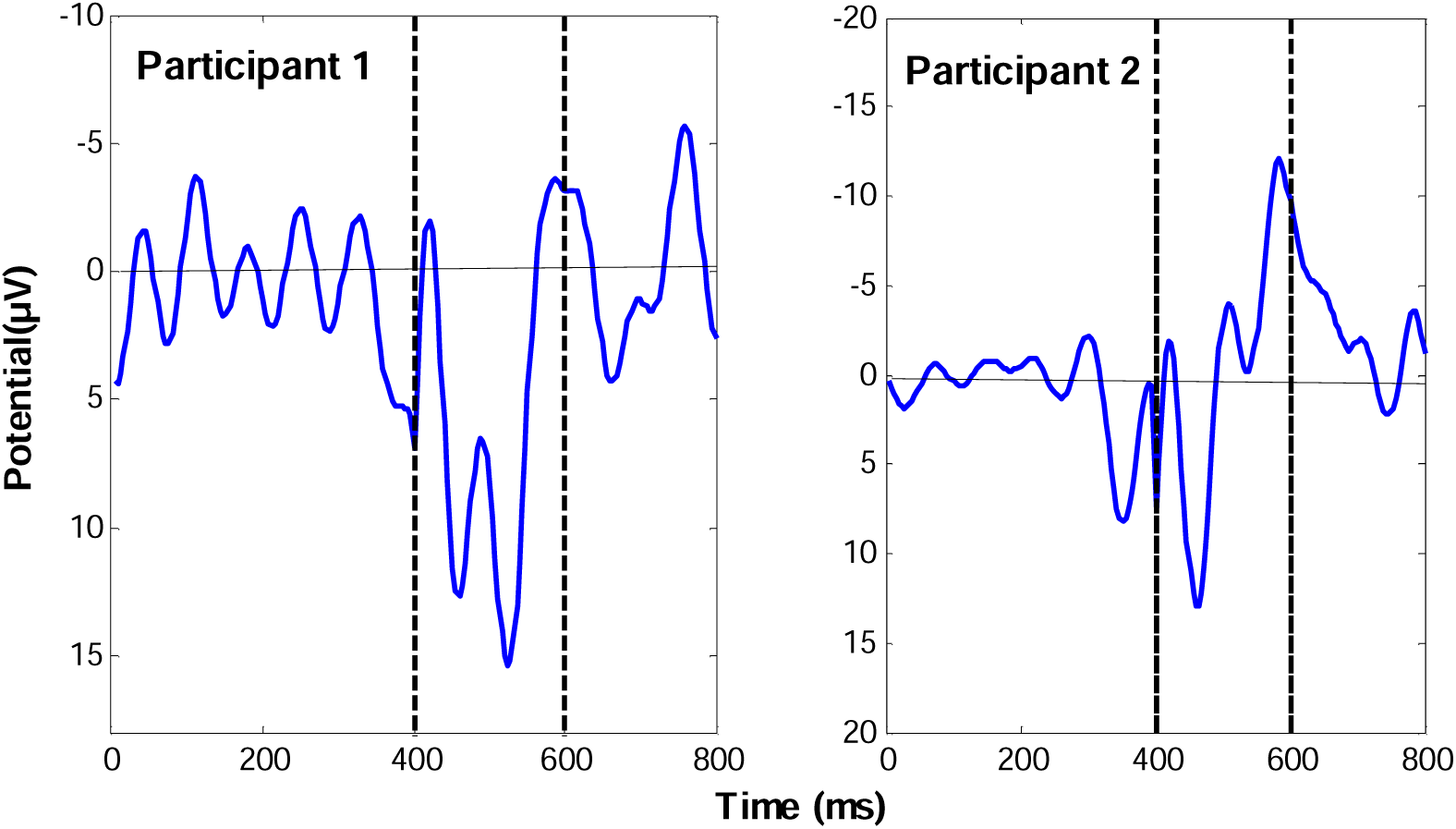
Measuring P3 size across participants using a fixed time window. When a fixed time window technique is used, each participant’s P3 is measured within the same time window. However, because of the expected variability across the participants, this procedure might yield false-negative cases. The Figure above, with positive plotted down, depicts an example of two individual ERPs. An a priori fixed window is placed from 400 to 600 ms. In such a situation, the P3 size of participant 1 will be measured appropriately with a large positive value because the P3 shape is almost always positive within the fixed time window. By contrast, although participant 2 has elicited a high positive P3 peak, the measure of the P3 size might result in a small positive value or even a zero or negative value due to the negative deflection that follows the positivity of the P3. In other words, within the fixed window, the classic negativity that follows the P3 positivity fell in the fixed window and cancelled that positivity. Such inter individual difference could arise for many reasons, e.g., participant alertness or variations in cortical folding.

In this paper, we explore the effectiveness of an ERP method that moves away from the fixed-form interpretation of components. Specifically, in a deception detection data set, we take advantage of a target detection condition, called the Fake in the experiment’s terminology, which is independent of the conditions that detect deception: the Probe and Irrelevant conditions. The critical point is that the ERP arising from the Fake accurately characterises the temporal profile elicited by each participant’s detection of a salient stimulus, i.e., it can be viewed as a *template* of such detection. This profile can then be used to test Probe vs. Irrelevant conditions for presence of such a pattern - guilt is determined by showing that the participant finds the Probe more salient than the Irrelevant. By analogy with *functional localisers* (Saxe, Brett, & Kanwisher, 2006), we call this a *functional profiler*. In fMRI, tasks are often included in experimental designs with the sole purpose of localising in space where a particular cognitive function manifests in participants’ brains, these are called functional localisers. The Fake ERP in this paper is conceptually used similarly but in respect of temporal profile, hence the term functional profiler.

Incorporation of such per-individual templating suggests adding hierarchy levels to the statistical model being applied. In typical group-level ERP analyses, a measure of interest is quantified for each participant; this might be area difference between two conditions, peak amplitude, latency, etc. A statistical test (t or ANOVA) is then run across these participant values. The quantification of error variance across participants introduces a distinct level, making this, strictly speaking, a hierarchical model with participants treated as a random effect (fixed effects would aggregate all participant single trials together for each condition, throwing away the participant distinction).

These traditional ERP models typically, though, maintain fixed-form components. That is, as previously discussed, identification parameters (such as window size and placement; and polarity) are fixed across participants, either by precedent from the literature or by inspection of the grand average (which is strictly fishing). If one is moving away from fixed-form components in order to accommodate inter-participant variance, it is natural also to consider identification parameters as a corresponding extra hierarchical level in the statistical model. For example, to accommodate inter participant variance, one might pick a different window or ERP template for each participant, which is legitimate fishing, as long as one formulates the test carefully in order to avoid type one error inflation, e.g., by using maximal statistics and/or permutation tests, as we do here.

In this context, this paper presents a methods-oriented demonstration of the increased statistical power arising from an application of a Monte Carlo permutation test, with (per-individual) Functional Profiling. We consider a number of identity deception detection data sets, in which we seek to find a differential ERP response for a Probe stimulus, i.e., the participants’ true birth date or true name. These data sets are taken from experiments demonstrating our Fringe-P3 deception detector (Bowman *et al.*, 2013, 2014) in which stimuli are presented so rapidly as to place them on the Fringe of awareness. The participant’s brain then searches for salient stimuli, performing so called (sub)liminal salience search, and participant detection of a salient stimulus, particularly the guilty knowledge, elicits a P3 ERP component. In this context, our decision to assess our analysis method on two data sets allows us to test it in both low (the birthdays data) and high (the names data) signal-to-noise situations. We also use simulated data to explore how the sensitivity and specificity of our method vary with parametric manipulation of the signal-to-noise ratio. At all stages, we verify that the type one error rate is not inflated relative to the alpha level. We explore the effectiveness of our methods both at a group and individual level.

### 1.1 Identity Deception Detection

P3-based deception detection systems employ the P3 component (also known as the P3) to detect concealed information. Several versions of these systems have been implemented, e.g., (Rosenfeld *et al.*, 2004; Abootalebi, Moradi and Khalilzadeh, 2006; Farwell, Richardson and Hernandez, 2006; Meixner and Rosenfeld, 2010a). Typically, these systems involve three main types of stimuli: Probes (**P**), which represent concealed information or crime details and can be recognized only by the guilty person; Irrelevants (**I**), which are frequent and task (crime)-irrelevant items; and Targets (**T**), which are similar to irrelevant items, but participants are asked beforehand to attend to these specific items and execute a task whenever they see one. For practical lie detection, the key comparison is between Probe and Irrelevant ERPs, since, for the *non-*deceiver, the former would be an Irrelevant. Importantly, the Probe for a deceiver typically generates a P3 ERP component, which is absent for the Irrelevant. Thus, the P3 Probe serves as an index of deception. The Target stimulus is presented for two reasons: first, it forces participants to pay attention to the presented stimuli, as they will be asked to do a task whenever they see a Target. That is, it shows whether a participant is cooperating or not. Second, as the Target stimuli are task-relevant items, they are expected to evoke a typical P3, which can be used in ERP assessment methods, such as the bootstrapped correlation difference method (Lawrence A. Farwell and Donchin, 1991). Note that, in our identity deception detectors, the term ‘Target’ was replaced by ‘Fake’, because in our paradigms participants were asked to detect their *Fake* birthdates and names. The following sections present the procedures of our identity deception detectors.

## 2 Methods

### 2.1 Birthday Deception Detection Experiment

The experiment presented in this section was introduced in (Filetti, 2013), in which the full experimental details are presented.

### 2.2 Names-Deception Detection Experiment

The experiment presented in this section was introduced in (Bowman *et al.*, 2013), in which the full experimental details are presented. Briefly, 15 participants were presented with 4 critical stimuli that we refer to as *Irrelevant1, Irrelevant2, Probe*, or *Fake.* These critical items could be a participant’s first name (Probe), their assumed name (Fake), or one of two preselected names unknown to the participant (Irrelevant1 or Irrelevant2). The concealed information was the participant’s first name (Probe). Critical items were presented in RSVP streams of name stimuli. Using a number of different P3 detection methods, e.g., our Weight Template (WT) and Peak-to-Peak, the aim was to investigate the difference in the brain responses of the participants to their first names (Probes) and to the irrelevant names (Irrelevant1 and Irrelevant2).

#### Innocents

As described in (Bowman *et al.*, 2013), the Name deception detector was also conducted on a control group (of Innocents) to assess the empirical false-positive rate. This Innocent group was treated exactly as the Guilty group, except that the participant’s real name was never presented. Thus, the Names identity detector was applied with 3 irrelevant names, giving 3 identical conditions: Irrelevant1, Irrelevant2, and Irrelevant3. These 3 Irrelevants yielded 6 pairwise comparisons because there are 6 permutations, e.g., (Irrelevant1, Irrelevant2), (Irrelevant1, Irrelevant3), (Irrelevant2, Irrelevant3), (Irrelevant2, Irrelevant1), and so on. We performed our statistical analysis on each such pair, with the first in the pair playing the (notional) Probe role and the second the Irrelevant role. Across the 8 participants who were assigned to the innocent group, there were 48 data sets, each comprising notional Probe and Irrelevant. We then analysed each data set with our ERP analysis methods, enabling a false alarm rate to be determined.

### 2.3 Methods for ERP Analyses

#### 2.3.1 Peak-to-Peak

In most P3-based lie detection studies, the Peak-to-Peak (i.e., amplitude difference) procedure is used to measure the disparity between deception detection ERPs (Rosenfeld, Ward, Frigo, et al. 2015; Labkovsky & J. Peter Rosenfeld 2012; Bowman et al. 2013). Several P3 concealed information test studies have shown that this procedure is more accurate and sensitive for detecting P3 deception compared to other methods such as baseline-peak and correlation difference (Rosenfeld, Ward, Frigo, et al. 2015; Soskins et al. 2001; Hu et al. 2012; Meixner et al. 2013). Thus, it was chosen to be compared to our proposed method: the WT approach.

The Peak-to-Peak measurement determines whether the amplitude of the P3 response elicited by a stimulus is greater than that elicited by another stimulus for the same individual. In line with (Rosenfeld et al. 2004; Labkovsky & J Peter Rosenfeld 2012; Rosenfeld, Ward, Frigo, et al. 2015), the word *peak*, in this paper, refers to averages across 100 ms intervals rather than single time points. In this study, this method was applied to measure the disparity between Probe and Irrelevant ERPs. First, subtracting the Irrelevant ERP from the Probe ERP generated a difference wave. This difference wave will contain a P3 pattern if the Probe condition elicited a bigger P3 response than the Irrelevant. To measure the P3, the Peak-to-Peak algorithm was used within a (defined) time window from 300 ms to 1000 ms from stimulus onset. In that time window, the algorithm searched for the maximal 100 ms interval (the highest peak). The algorithm then searched from the first time point that follows this maximal 100 ms interval until the end of the defined time window for the minimum 100 ms interval (the lowest peak). The difference between the highest and lowest peaks was then calculated to define the Peak-to-Peak measure. To evaluate the statistical significance of this measure, a randomisation (i.e., Monte Carlo permutation) test was applied to generate a null hypothesis distribution. From this distribution, a p-value can be inferred. If the p-value < .05, then the null hypothesis that the Probe and Irrelevant ERPs are the same is rejected (for additional details see the Monte Carlo permutation test section).

#### 2.3.2 Weight Template (WT)

The new method presented here uses an individual-specific template to search for the P3 response. This template characterises an individual’s P3 shape from the Fake condition, where the largest P3 is expected; remember, the key guilt-determining comparison is between Probe and Irrelevant, with the Fake typically being ignored. This WT approach handles situations in which the shape of an individual’s P3 is atypical. The *Template* was defined from the difference between the Fake and Irrelevant ERPs. This template was then multiplied (point-wise) by the difference between the Probe and Irrelevant ERPs. The result of this multiplication was used to determine whether a participant was guilty or not. In more detail, firstly each ERP was normalised, resulting in the mean over time being zero and the standard deviation being one (i.e., each ERP point becomes a Z-score) (Bandt *et al.*, 2009).

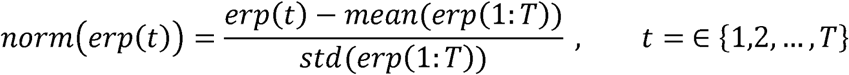

Where *erp*(*t*) is a single ERP that is a function of time (*t*),*std*(*erp*(1:*T*)) is the standard deviation and *mean*(*erp*(1:*T*)) the mean across the ERP’s time points and *T* is the number of time points in the ERP. Secondly, once Probe, Irrelevant, and Fake ERPs were normalised, the point-wise difference between the normalised Fake ERP *norm*(*F*) and the normalised Irrelevant ERP, *norm*(*I*), was determined. The result was a vector that we refer to as the *weightVector* :

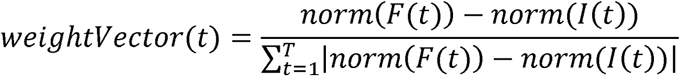

The *weightVector* was used as a template, with which the difference between the normalised Probe ERP *norm*(*P*) and the normalised Irrelevant ERP *norm*(*I*) was multiplied:

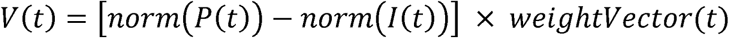

Finally, the P3 weighted difference was obtained by calculating the sum of *V* (*t*) over time :

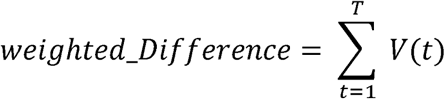

For the purpose of this study, we used the weighted difference value as a scale for the presence or absence of the P3 in the Probe ERP. The larger this value is, the more likely the P3 was elicited and the more likely a guilty decision could be made. To evaluate this value statistically, a randomisation (i.e., Monte Carlo permutation) test was applied. Thus, for each participant, a null hypothesis distribution was generated, and a p-value was then calculated. To reject the null hypothesis that Probe and Irrelevant ERPs are not different, and thereby detect a participant as guilty, the p-value should be less than 0.05.

### 2.4 Statistical analyses

#### 2.4.1 Monte Carlo Permutation Test

As mentioned in the previous section, the performance of the methods was evaluated using Monte Carlo permutation resampling (randomisation test), which has been extensively used to analyse EEG (Blair and Karniski, 1993; Groppe, Urbach and Kutas, 2011) and fMRI data (Brammer *et al.*, 1997). As a central purpose of this analysis was to classify participants into guilty and innocent, the methods were applied on a per-participant basis (individual-level analysis). In general, a randomisation test involves the following two steps: (a) the computation of some statistic on the true observed data and (b) an evaluation of its p-value under the randomisation distribution. This distribution arises as a result of the random assignment of the units (in our case, *trials*) to conditions.

Because the key comparison in our ERP analysis methods is between the Probe and Irrelevant conditions, we applied the randomization test in the same manner as it used in (Bowman *et al.*, 2014).

#### 2.4.2 Group and Individual-Level Analysis

We assessed our methods on both a per-individual, which is required in many applications, including P3-based concealed Information Test, and on a group-level basis, as required for a population-level inference. At the individual-level, we determined whether the brain response evoked by the Probe stimulus was significantly different from that evoked by the Irrelevant within an individual. As described earlier, a randomisation was used to generate a p-value for each participant.

To determine whether there was a significant difference between the Probe and Irrelevant responses at the group level, a paired t-test was used. Each participant was assigned a pair of Peak-to-Peak sizes, one from Probe and one from Irrelevant. A paired t-test was then applied across participants in each experiment. Peak-to-Peak sizes of both conditions in Names and Birthdays data sets are presented in tables 1 and 2 respectively.

**Table 1.** Grand average of p-values at different levels of SNR. presents the grand average p-values and SNRs for each method across all participants. For each K (K was used as a scalar for the amplitude of noise), the grand average values were obtained by taking the average of all p-values and SNRs that are presented in Table A.1 in Appendix A. The confidence intervals indicate the 95% confidence intervals for the means as estimated by bootstrapping over p-values (see section 3.1.1 for more details about estimating the 95% confidence interval). As *K* increases, the SNR declines, and therefore the p-value increases. The two methods performed almost equally at a high SNR. However, at low SNRs, the p-values of the WT were significantly smaller than the corresponding values for Peak-to-Peak.

| K | SNR | WT |  |  | Peak-to-Peak |  |  |
| --- | --- | --- | --- | --- | --- | --- | --- |
|  |  | Grand average | 95% Ci |  | Grand average p value | 95% Ci |  |
|  |  | P value |  |  |  |  |  |
| 1 | .1360 | <.0001 | .0000 | .0000 | <.0001 | .0000 | .0000 |
| 7 | .0404 | .0006 | .0004 | .0011 | .0006 | .0003 | .0012 |
| 13 | .0163 | .0099 | .0078 | .0128 | .0182 | .0151 | .0226 |
| 19 | .0086 | .0277 | .0238 | .0329 | .0503 | .0442 | .0581 |
| 25 | .0052 | .0641 | .0576 | .0712 | .1058 | .0968 | .1158 |
| 31 | .0035 | .1010 | .0930 | .1099 | .1468 | .1359 | .1591 |
| 37 | .0025 | .1301 | .1209 | .1400 | .1827 | .1704 | .1954 |
| 43 | .0018 | .1620 | .1513 | .1727 | .2107 | .1978 | .2239 |
| 49 | .0014 | .1925 | .1817 | .2044 | .2531 | .2398 | .2671 |

**Table 2.** Grand average of AUC versus SNR. presents the grand average AUCs and SNRs for each method across all participants in in the Names experiment. For each noise level (K), the grand average values were obtained by taking the average of all AUCs and SNRs that are presented in Table A.2 in Appendix A. It also shows results of paired t-tests (t and p values) that were applied to AUCs at each SNR level. Section 3.1.2 describes how the t-test was applied at each SNR level. As can be seen, the average performance of both methods decreased as SNR decreased. However, as the SNR dropped, the difference in average performance of the WT compared to Peak-to-Peak increased. Results of a paired t-test at each SNR level statistically confirmed this pattern.

| K | SNR | Grand average of AUCs |  | Paired t-test |  |
| --- | --- | --- | --- | --- | --- |
|  |  | WT | Peak-to-Peak | t value | P value |
| 1 | .1360 | 1.000 | 1.000 | NaN | NaN |
| 7 | .0404 | .999 | .999 | -.5405 | .5973 |
| 13 | .0163 | .988 | .982 | .9520 | .3572 |
| 19 | .0086 | .969 | .950 | 1.6401 | .1233 |
| 25 | .0052 | .929 | .895 | 2.0528 | .0593 |
| 31 | .0035 | .889 | .854 | 2.3244 | .0357 |
| 37 | .0025 | .857 | .818 | 2.7291 | .0163 |
| 43 | .0018 | .822 | .790 | 2.9170 | .0113 |
| 49 | .0014 | .788 | .746 | 3.4875 | .0036 |

In the WT method, we compared Probe and Irrelevant conditions by generating WT values per-participant for each condition. The Fake served as a template and was applied as described earlier except only the Fake condition was used, i.e.,

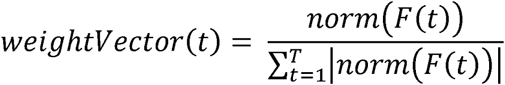

Then, for example to obtain a WT value of condition X, we applied the *weightVector*(*t*) to X as,

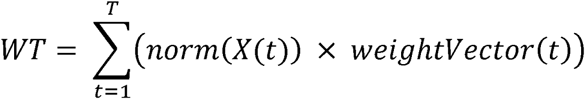

A paired t-test was then applied to the WT values of both conditions across all participants. The WT values of both conditions in the Names and Birthdays data sets are presented in Tables 2.

### 2.5 Simulated EEG Data

The aim of generating simulated data was to explore how the sensitivity and specificity of the WT method vary with parametric manipulation of the Signal-to-Noise Ratio (SNR).

#### 2.5.1 Intrinsic validity of the WT

To explore the validity of the WT method, we applied it to a set of artificial data for which we already know that the null hypothesis is true (i.e., there is no significant difference between the conditions). The aim was to assess the false-positive rate (i.e., the Type I error rate) of the method. Statistically, false-positive errors occur when a statistical test rejects the null hypothesis (H0) when it is in fact true (i.e., the WT indicates a significant difference between two conditions when none is present). The probability of a Type I error is always set in advance and should be equal to the alpha level of a statistical test, in our case .05.

We assessed our method in two intrinsic validity tests: the first was on pure noise data, while the second was on real human ERPs plus random noise. In the first test, we generated a set of simulated EEG data using the function *noise* that was used in (Yeung *et al.*, 2004, 2007); this contains auto-correlation statistics consistent with human EEG^1^. Specifically, we generated 1000 simulated EEG data sets. Each data set was composed of 150 random noise trials. These trials were then randomly split into three equal sets, which we refer to as simulated Fake, Probe, and Irrelevant conditions; thus, each condition comprised 50 trials. A simulated ERP was then generated for each condition by taking the average of the 50 trials. Using our Monte Carlo permutation test, we applied the WT method to each data set in an identical manner as in our analyses of the real experimental data. In this way, applying the WT to each data set will yield 1000 p-values. The Type I error rate was then defined as the proportion of p-values below .05.

In the second validity test, we assessed the Type I error of the method in a slightly different way, in which we compared between two conditions that contain a real human ERP signal. Particularly, we explore the validity of the WT approach by creating two artificial conditions (Probe and Irrelevant) that have the same signal in, where *signal* refers to an ERP containing a P3. The reason for doing such an assesment is that the Irrelevant ERP may have a similar shape to a Probe (Bruno Verschuere, Gershon Ben-Shakhar, 2011). In other words, in some cases, an Irrelevant ERP may contain a signal (a small P3) as can be seen in (Lawrence A. Farwell and Donchin, 1991; Rosenfeld *et al.*, 2004). Indeed in general, ERPs may contain some sort of response to a stimulus, which is a signal, and the null hypothesis is that the form of this signal does not change across conditions. Thus, it was vital to assess the Type I error rate of our method when conditions have similar shapes. The procedure was as follows: we created two artifitial conditions (Probe and Irrelevant), using the same signal (human ERP) plus pure random noise. This signal was in fact the grand average of the Probe condition in the Names experiment (see Figure 2). As in the first test, we again generated 1000 simulated EEG data sets, each consisting of 50 Probe trials and 50 Irrelevant trials, in which each trial of both conditions contained the same signal added to random noise, that is,

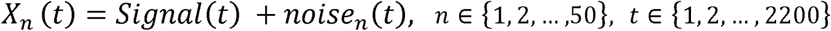

where *t* is a time point and *n* a trial. An artificial ERP was then generated for each condition by taking the average of the 50 trials. We then applied the WT method to each condition in exactly the way we did in our analyses of the real experimental data. In particular, the *Template* for the WT method was constructed using the grand average of the Fake condition in the Names experiment (see Figure 2). In this way, this test will yield 1000 p-values. The Type I error rate was then defined as the proportion of p-values below .05. In these first and second tests, the parameters used to generate the noise time series were exactly the same; the length of a single trial was 2200 time points and the sampling frequency was 2000 Hz.

**Figure 2.**
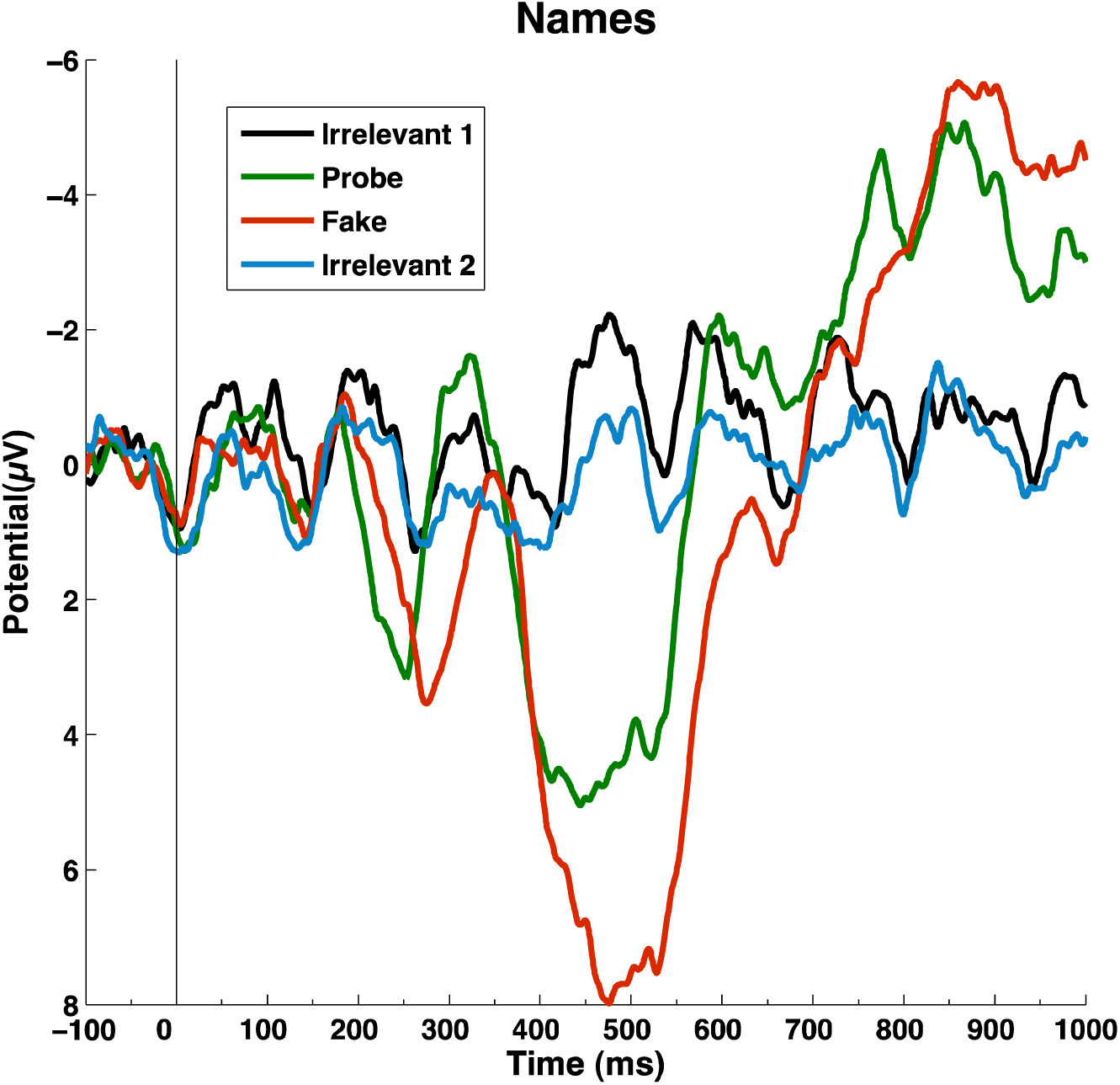
Pz grand averages for Names experiment. The grand averages at Pz for all of the conditions in the Names deception detector. Positive is plotted downwards.

#### 2.5.2 Comparing method’s performance at different level of SNR

The aim of generating simulated data was to evaluate the performance of the WT and Peak-to-Peak methods with different levels of SNR (Signal-to-Noise Ratio). We varied the SNR by adding a different amount of noise to the Probe and Irrelevant ERPs of each participant in the Names deception detector experiment. Specifically, first, Probe and Irrelevant ERPs were generated in the normal manner. Second, we generated a set of *Surrogate* trials, each of which was the addition of a condition ERP and a single noise trial. For example, a single *surrogate* Probe trial can be defined as the following:

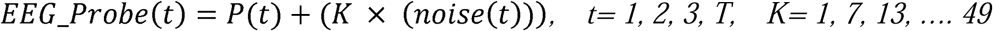

Where *P* is the participant’s Probe ERP, *noise* is a vector of (auto-correlated) random simulated noise, and *T* is the number of time points. To vary the amount of noise in a single trial, we scaled the amplitude of the *noise* by multiplying the vector that represents it by a number (a ‘scalar’), which we denote as *K*. In this way, a surrogate Probe ERP was created by generating (*N* =50) *surrogate* Probe trials and then taking the average of these N *surrogate* trials. A surrogate Irrelevant was created in the same way. Third, we estimated the SNR of the surrogate Probe and Irrelevant ERPs by calculating the ratio between the root mean square of the signal (Probe/Irrelevant) and the *noise* (estimating the SNR process will be explained in the next section). Next, we applied a randomisation test with our main methods (Peak-to-Peak or WT) to both surrogate ERPs providing p-values in an identical manner as with the real experimental data (see the randomisation test section). 100 iterations of these steps will yield 100 p-values for each method and 100 SNR estimates for each condition. Next, a single average p-value (across the 100 individual p-values) was obtained for each method. In addition, for each surrogate condition, an average *SNR*_*avg*_ value was estimated (across the 100 SNRs). This procedure was repeated for every *K* value (from 1 to 49), in steps of 6.

This procedure evaluates the performance of each method in 2 different ways. First, the procedure compares the average of 100 p-values of each method at different levels of SNR; the smaller an average p-value relative to a particular alpha level (in our case .05), the greater the statistical power of that method. Second, the procedure calculates the area under a ROC (Receiver Operating Characteristic) curve (abbreviated AUC) for each method; see next section. The above procedure can be summarised as follows:

i. For each participant, determine the Probe and Irrelevant ERPs.
ii. Set K=1
iii. For a given K, generate surrogate (noisy) Probe and Irrelevant ERPs, each of which is composed of 50 trials.
iv. Estimate the SNR of the surrogate Probe and Irrelevant.
v. Apply a randomisation test with Peak-to-Peak and WT methods to the surrogate ERPs, thereby generate a null hypothesis distribution and calculate a p-value for each method.
vi. Repeat the previous steps (iii-v) 100 times.
vii. For each method, calculate the average of the 100 p-values that were obtained by the 100 iterations as well as the average SNR (*SNR*_*avg*_) for each condition.
viii. For *K* <= 49, increase *K* by 6 and return to step iii. ix.

#### 2.5.3 Statistical Analyses on the Simulated Data

##### Signal-to-Noise Ratio (SNR)

In our simulated data analyses, the SNR was used to quantify the ratio between the ERPs (Probe/Irrelevant) and the noise. For each condition, the SNR was estimated from the ratio between the root mean square (RMS) amplitudes of an ERP (Probe/ Irrelevant) and the noise (Drongelen, 2006). First, for each single trial *n*, we estimated the SNR as follows:

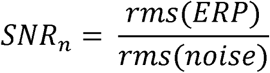

where the *rms* of time series *x* across time points *t* can be defined as (van Drongelen, 2006),

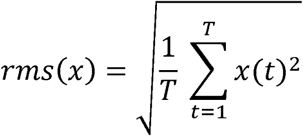

The *rms* of an ERP was calculated in a time window from stimulus onset to 1000 ms post-stimulus. We calculated the amount of noise in the ERP level as

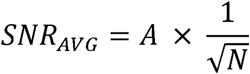

where *A* is the average of all of the SNRs across *N* trials, i.e.,

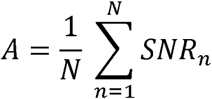

##### ROC curve

A receiver operating characteristic (ROC) curve is a means to evaluate the performance of classifiers as their discrimination criterion (threshold) is varied. ROC graphs are used in signal detection theory to depict the trade-off between the sensitivity (hit rate) and specificity (false alarm rates) of classifiers(Egan, 1975; Van Erkel and Pattynama, 1998). A ROC graph is a 2-dimensional graph in which the true-positive (hit) rate is plotted on the Y axis and the false positive rate is plotted on the X axis. A single point in ROC space represents an output of a classifier in terms of the hit (true positive) rate and the false-positive rate. To quantify ROC performance, measurement of the area under the ROC curve (AUC) is used *(Fawcett, 2006).* In this study, ROC curve analysis was used to evaluate the performance of our methods with simulated data. In our simulated data, for each SNR value, 100 p-values were obtained by each method. For each method, we then calculated the true-positive rate as the number of p-values that were less than a certain alpha level threshold divided by 100. To calculate the false-positive rate, we created 2 data sets that were composed of (pure) noise trials and that did not contain any signal. Specifically, we used the same noise function to generate at random 100 noisy trials; the average of the first 50 trials played the role of a Probe and the average of the remaining trials played the role of an Irrelevant. We then applied our methods with a randomisation test exactly as described earlier. Here, 100 iterations of this process will yield 100 p-values. The false-positive rate was then calculated at different alpha levels as the number of p-values that were less than that alpha level divided by 100. The threshold (alpha level) varied from 0 to 1, with a step of .01, to estimate both rates.

## 3 Results

### 3.1 Results of Simulated Data Analysis

As the WT method is new, we must first confirm its validity. Thus, the Type I error rate, ‘the true false-positive rate’, of the method was assessed on simulated data. Statistically, the Type I error rate of the method should be .05 (our alpha level). As described earlier, we assessed this rate in two tests: in the first test, we generated 1,000 groups of artificial data sets (pure noise) for all of the conditions (Fake, Probe, and Irrelevant). In the second test, we compared between two artificial conditions (Probe and Irrelevant) with the same signal. The average p-values of 1,000 WT runs in the first and second tests were .489 and .504, respectively. The average p-value of both tests was indeed approximately .5, as expected when there is no difference between the Probe and the Irrelevant conditions. Importantly, the Type I error rates of the method in the first and second tests were .039 and .0484. These results confirm that the false-positive rate of the method is not larger than our alpha level of .05. Figure 4 shows the uniform distributions of the 1,000 p-values for both tests.

**Figure 3.**
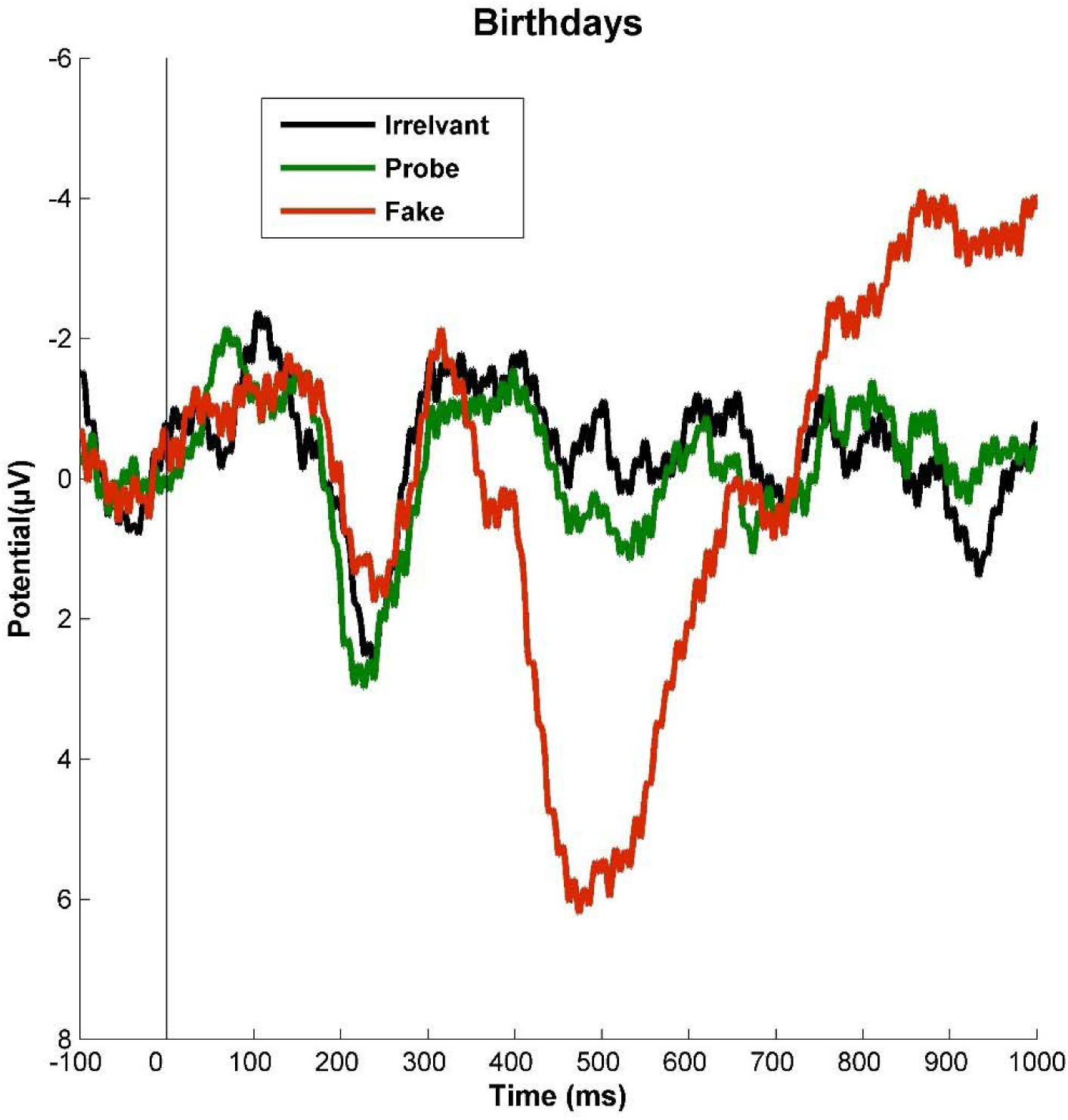
Pz grand averages ERPs for Birthdays experiment. The grand averages at Pz for all of the conditions in the Birthdays deception detector. Positive is plotted downwards.

**Figure 4.**
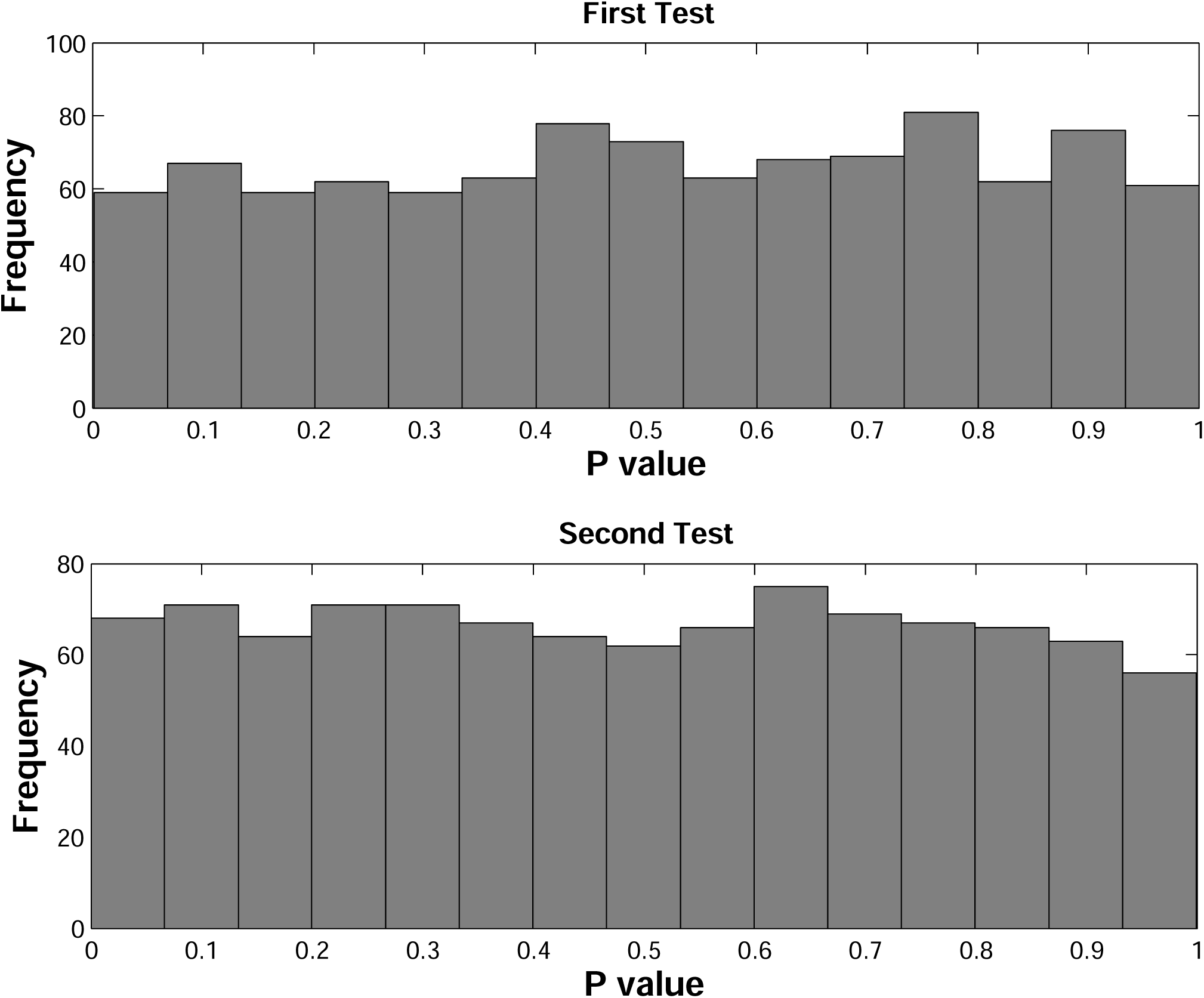
P-value distributions obtained from the intrinsic validity test of the WT. The upper and lower plots show the distributions of p-values that were generated from the first and second intrinsic validity test. Each distribution consists of 1000 p-values. As expected, both distributions are uniform.

#### 3.1.1 P-value by SNR

As noted previously, the aim of generating artificial data sets was to evaluate the WT methods on data with pre-defined settings, such as the SNR level. Specifically, we were interested in comparing the performance of the WT and Peak-to-Peak with varying SNRs.

The statistical analyses showed that the performance of both methods decreased as the SNR declined, as expected. As the amount of noise increased, the p-values increased towards .5; the p-value corresponding to identical Probe and Irrelevant. At very high SNRs, both of the methods performed successfully, with nearly equal p-values across all participants. For the participants who elicited sharp and well-defined Peaks, both methods continued to detect the peaks successfully, even at low SNR levels. For example, participant 8 (see Figure 4) has high positive P3 peaks, which yielded significant p-values (p<.001), even at low SNRs of .02. As SNRs dropped, the WT displayed increasingly better performance relative to Peak-to-Peak, with the p-values of the WT generally smaller than the corresponding p-value of the Peak-to-Peak.

The per-individual p-values at each SNR level for both methods are presented in Table A.1 in Appendix A. At each SNR, the grand average p-values for both methods were obtained by taking the mean of all p-values across all participants. This was done at each SNR. Table 1 showed that at high SNRs, the grand mean p-values of both methods were nearly the same whereas, at low SNRs, the grand mean p-values of the WT were less than the grand mean p-values of Peak-to-Peak. Moreover, in order to obtain confidence intervals, a non-parametric bootstrap confidence interval method was used (Wasserman and Bockenholt, 1989; Carpenter and Bithell, 2000). Specifically, for each SNR, 95% confidence intervals for the grand average p-value were obtained by bootstrapping 1000 times over 1500 p-values (15 participants × 100 p-values). The grand average p-values and 95% confidence intervals are shown in Table 1. Figure 6 shows the grand average p-value as a function of the SNR. As can be seen in Figure 6, as SNR decreased, the distance between confidence intervals of the two methods increased suggesting an increasing (statistically significant) difference between methods.

**Figure 5.**
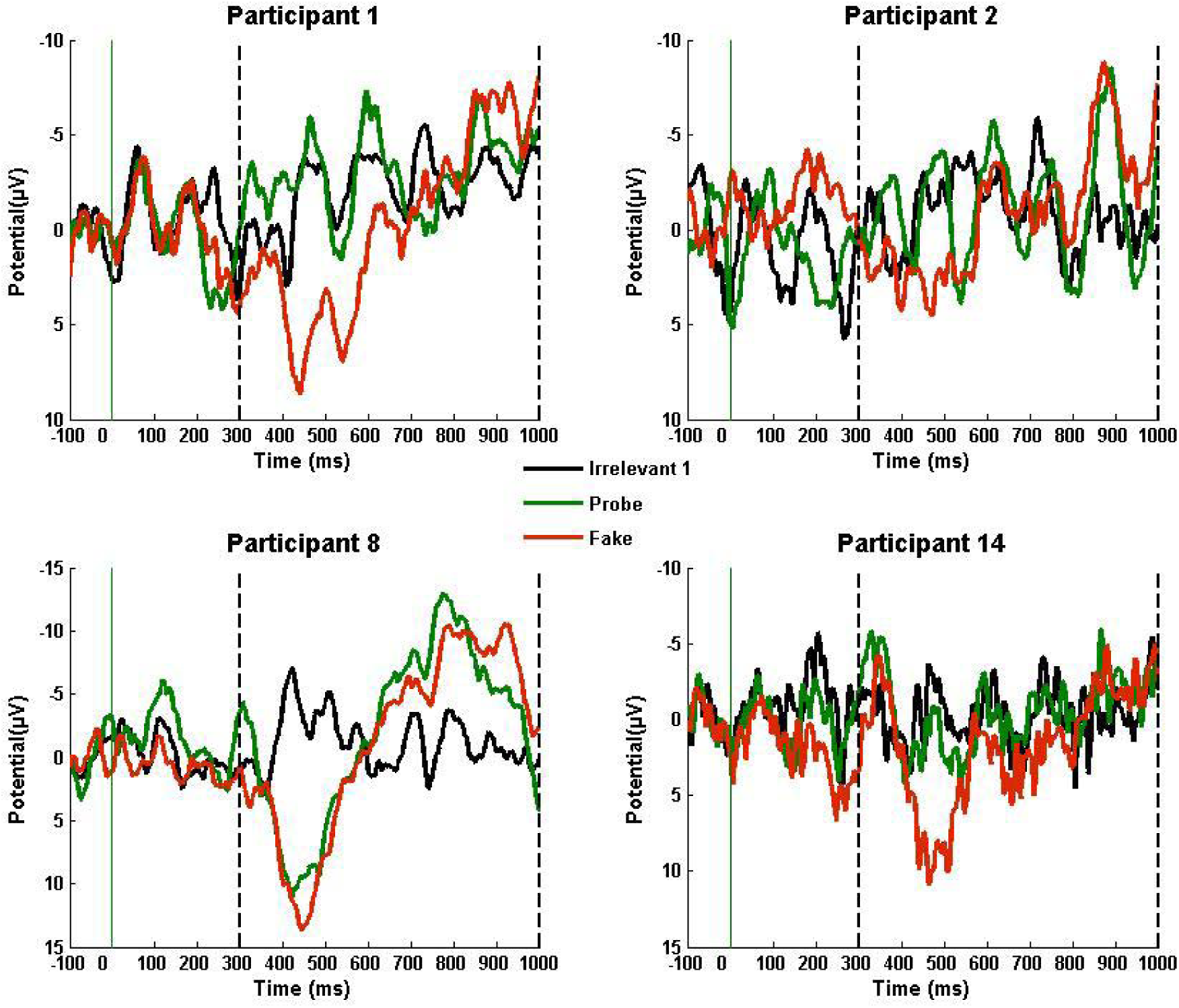
Selection of Pz ERPs form Names Experiment. Selections of individual ERPs at Pz for participants 1, 2, 8, and 14 in the Names experiment. Positive is down, and the vertical dashed lines mark the region in which we search for the P3.

**Figure 6.**
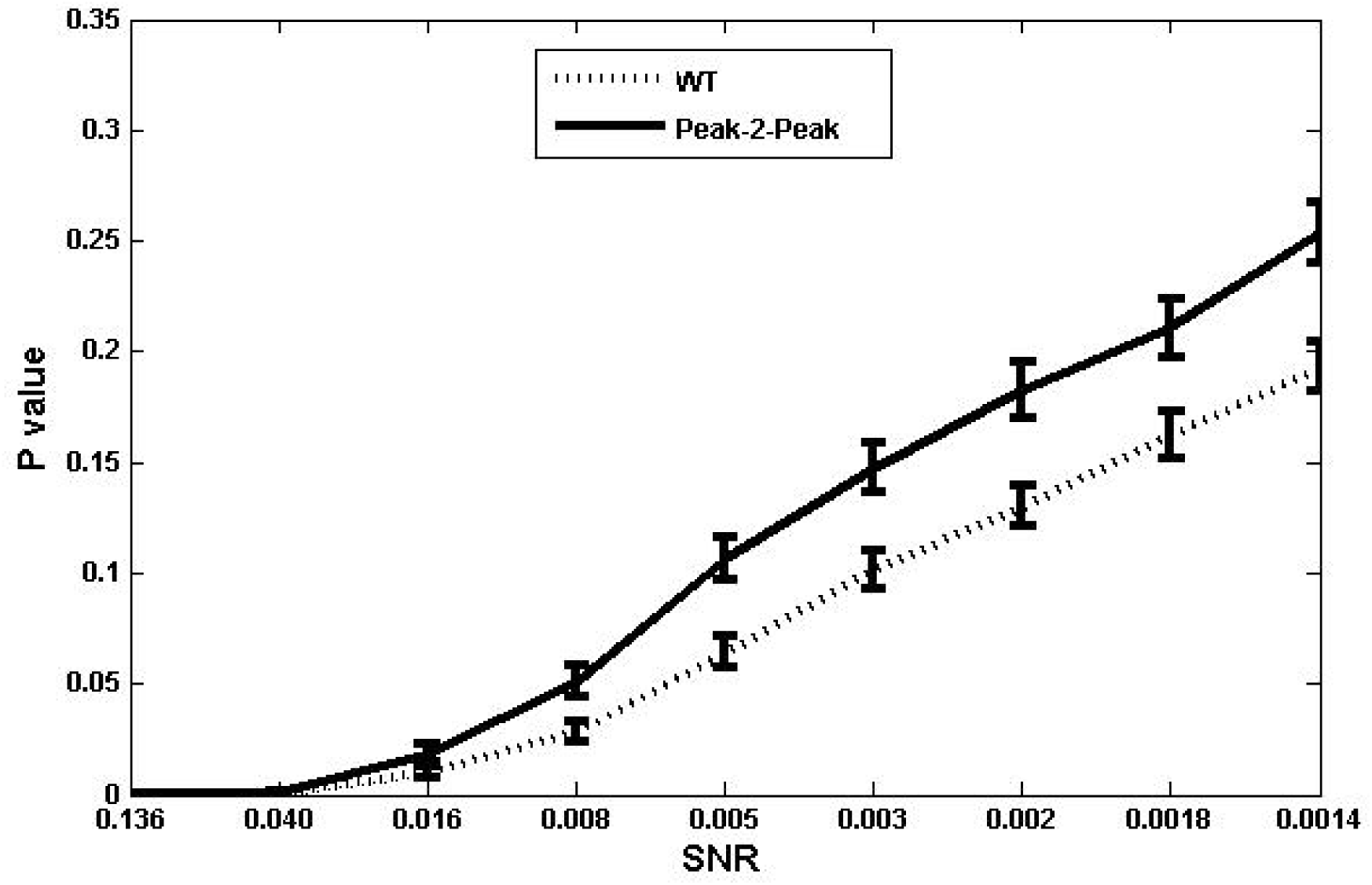
Grand average p-values at different level of SNR. The figure shows the grand average p-value as a function of the grand average SNR. At each SNR level, the grand average p-value was obtained by taking the mean of all p-values presented in table A.1. For example, the grand average p-value at a certain SNR level was calculated by taking the average of all 15 (across all participants) p-values that were obtained at that level. The Grand average SNR was calculated in the same way by taking the average of all estimated SNRs. As shown in the figure, the p-values of both methods increase as the amount of noise increases (the SNR decreases). As the SNR decreased, the amount p-values of the WT were smaller than the corresponding p-values of the Peak-to-Peak increased. Error bars indicate the 95% confidence intervals for the means as estimated by bootstrapping over p-values at each SNR.

#### 3.1.2 AUC by SNR

Overall, our results also indicate that, broadly speaking, with high SNR ERPs, the two methods are equally successful at detecting deception, i.e., the P3 component. However, at low SNRs, the WT significantly outperforms the Peak-to-Peak. As can be noted from Table A.2, as the K value increased, SNR decreased towards 0, and therefore the AUC of both methods decreased towards .5. The AUC measures of both methods demonstrated that for the participants who elicited a clear and high P3 component, the performance of the two methods were almost equal, reaching towards AUC=1. For example, as shown in Figure 4, participant 8 has a large positive P3 peak (amplitude>10 µv) and, although a large amount of noise was added (K <= 31) to the Probe waveform, the P3 peak remained stable and resistant, and thus both of the classifiers detected the component successfully with AUC>.99. However, for different levels of SNR across all individuals, the performance of the WT was more resistant to noise. The lowest AUC for the WT was approximately .54, for participant 1, and this result was not surprising since before noise was added, there was already considerable noise in the Probe waveform, and there was no evidence of P3 presence (Figure 4 depicts ERPs of participants 1 in the Names experiment). Figure 7 shows selected ROC curves considering different levels of SNR (different K values). The ROC curves are presented at three different K values (K=13, 31, and 49) of the WT and Peak-to-Peak for the simulated data from participants 1, 2 and14. For each participant, as K increases (i.e., the noise increases), the ROC curves of both methods move from the upper left corner towards the diagonal (i.e., the performance of each classifier decreases).

**Figure 7.**
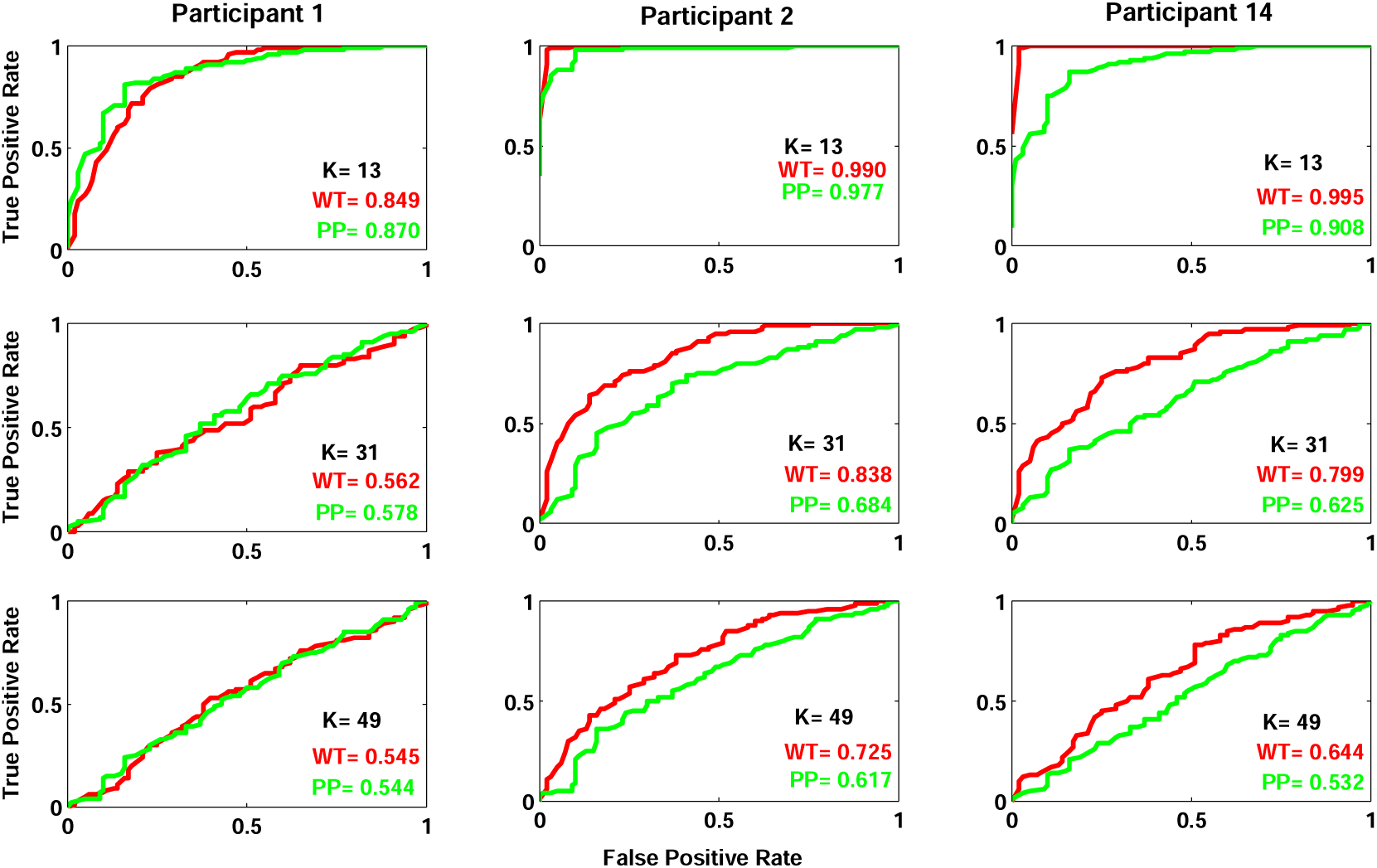
Selection of ROC curves. The ROC curves of the WT (red) and Peak-to-Peak (green) methods, using simulated data for the ERPs of participants 1, 2, and 14. Each column in the figure shows the performance of each classifier for each participant at different noise levels (K), K= 13, 31, and 49. The ROC curve depicts the relationship between the false-positive rate (FPR) (X axis) and the true-positive Rate (TPR) (Y axis). Note the K is used as a scalar for the amplitude of noise.

The per-individual AUC scores at different levels of SNR are presented in Table A.2 in Appendix A. Furthermore, in order to confirm the statistical difference between performances of the two methods at a group level, a paired t-test was applied to AUC scores at each SNR level across all 15 participants. That is, a paired t-test at each SNR level was applied to the 15 (no. of participant) AUC pairs (one for each method) to infer whether the performance of methods are statistically different. Table 2 shows the grand average AUC scores and the outcomes of the paired t-tests, which show that there was no significant difference between the performances of methods at high SNR levels. However, there was a significant (and increasing) difference as the SNRs decreased.

### 3.2 Results of Experimental Data

#### 3.2.1 ERPs of the Birthadays and Names data sets

Because the P3 amplitude is expected to be maximal over the parietal areas of the scalp (Lawrence A. Farwell and Donchin, 1991), our analysis was focused on the Pz channel. This is also in line with (Labkovsky & Rosenfeld 2012; Rosenfeld et al. 2015). Figure 2 shows the grand average ERPs for the 4 conditions in the Names experiment. Large P3 responses were elicited by Probe and Fake but not by Irrelevant1 or Irrelevant2. Irelevant1 and Irrelevant2 are very similar.

Figure 3 shows the grand average ERPs (across all of the participants) of the Birthdays experiment in the Guilty group for the Fake, Probe, and Irrelevant responses. The Irrelevant and Probe patterns are similar, with the exception that, in the typical P3 time window, the Probe elicited a small P3, whereas the Fake pattern significantly differed from the other two responses, showing a large and clear P3.

We performed WT and Peak-to-Peak measurements at a single subject level for measuring the statistical significance of the brain responses elicited by the Probe and Irrelevant conditions. As noted, to diagnose guilt or innocence, the P3 responses of both conditions were compared: in a Guilty participant, Probe is expected to be larger than Irrelevant; in an Innocent participant, no difference is expected between the conditions because the Probe is just another Irrelevant. A randomisation technique was used with each analysis method, Peak-to-Peak and WT, generating a p-value for each participant. In the Names Experiment, since we have two Irrelevants (Irrelevant1 and Irrelevant2) conditions, the key comparison was between Probe and Irrelevant1. All classification results were calculated at an alpha level of .05.

#### 3.2.2 Hit and false alarm rates

In the Names experiment, with the peak-to-peak method, 12 participants (12/15) in the Guilty group were detected as guilty (80% hit rate). The false alarm rate using Peak-to-Peak was 8% (4/48). In the Birthdays experiment, the hit rate was 17% (2/12), and the false alarm rate was 20% (2/10). The WT had a hit rate of 93% in the Names experiment; 14 participants (14/15) in the Guilty group were detected as guilty, while the false alarm rate was 2% (1/48). In the Birthdays experiment, the hit rate was slightly better than for the peak-to-peak, 33% (4/12), with a false alarm rate of 10% (1/10). Tables 3 and 4 present per-individual p-values arising from both methods for the Birthdays and Names experiments, respectively. The false-positive test outcomes for the Names and Birthdays data sets are shown in Tables 5 and 6 respectively.

**Table 3.** Per-individual analyses for the Names experiment. This table presents true observed values and p-values obtained from the WT and Peak-to-Peak methods for each participant in the Names experiment. The WT values of the Probe and Irrelevant1 conditions were obtained by applying the Fake as a template to each condition. The Peak-to-Peak method was applied to each condition (Probe and Irrelevant1) to obtain their Peak-to-Peak sizes. At an alpha level of .05, 14/15 of the participants were detected as guilty with the WT, whereas 12/15 of the participants were found to be guilty using Peak-to-Peak. Note that he smallest p-value we can obtain with a 1000 resamplings is 0.001. However, the exact p-values for many of the participants are likely to be significantly smaller than <0.001.

| Participant | WT |  |  | Peak-to-Peak |  |  |
| --- | --- | --- | --- | --- | --- | --- |
|  | Probe | Irrelevant1 | P value | Probe | Irrelevant1 | P value |
| 1 | .4855 | .4374 | .342 | 4.2433 | 4.2342 | .286 |
| 2 | .5705 | -.0925 | .011 | 4.2141 | 3.8075 | .080 |
| 3 | .9828 | .1141 | <.001 | 11.0444 | 3.0311 | <.001 |
| 4 | .8973 | -.1847 | <.001 | 7.8579 | 4.2099 | <.001 |
| 5 | .9184 | -.0159 | <.001 | 13.6103 | 2.8520 | <.001 |
| 6 | .9921 | .4822 | <.001 | 26.224 | 2.8163 | <.001 |
| 7 | .9466 | -.2069 | <.001 | 11.629 | 0.9923 | <.001 |
| 8 | 1.0784 | -.4384 | <.001 | 20.406 | 2.2978 | <.001 |
| 9 | 1.0085 | .3274 | <.001 | 10.643 | 3.1951 | <.001 |
| 10 | .6516 | -.1132 | .001 | 9.957 | 5.3638 | .001 |
| 11 | .4589 | -.2025 | .019 | 6.0047 | 2.2418 | .023 |
| 12 | .9898 | .3093 | <.001 | 15.6686 | 2.3618 | <.001 |
| 13 | .6292 | -.4560 | <.001 | 11.5548 | 2.7867 | <.001 |
| 14 | .8114 | .1305 | .012 | 4.4183 | 2.5450 | .239 |
| 15 | 1.0423 | .3863 | <.001 | 9.9656 | 2.3088 | <.001 |

**Table 4.** Per-individual p-values for the Birthdays experiment. shows true observed values and p-values arising from the WT and Peak-to-Peak methods for each participant in the Birthdays experiment. At an alpha level of .05, 4/12 of the participants were detected as guilty with the WT, whereas 2/12 of the participants were found guilty using Peak-to-Peak.

| Participant | WT |  |  | Peak-to-Peak |  |  |
| --- | --- | --- | --- | --- | --- | --- |
|  | Probe | Irrelevant | P values | Probe | Irrelevant | P values |
| 1 | .9553 | .6907 | .0760 | 7.4228 | 4.1333 | .080 |
| 2 | .3772 | .7839 | .4780 | 2.6645 | 3.8197 | .419 |
| 3 | .6277 | .0031 | .0480 | 6.1961 | .9474 | .119 |
| 4 | -.1725 | .2530 | .9540 | 3.5902 | 5.9752 | .478 |
| 5 | .5486 | -.1014 | .0540 | 4.2658 | 4.6447 | .125 |
| 6 | .6776 | -.2415 | .0140 | 4.9793 | 3.3458 | .040 |
| 7 | -.0933 | -.6693 | .0200 | 4.0758 | 3.1902 | .086 |
| 8 | .2194 | -.1179 | .2290 | 2.5920 | 2.7268 | .536 |
| 9 | .3813 | .0215 | .1850 | 2.1699 | 1.8156 | .312 |
| 10 | .3108 | .5748 | .8700 | 3.3671 | 4.2875 | .988 |
| 11 | 1.0487 | .9481 | .5090 | 5.5972 | 3.3216 | .414 |
| 12 | .4629 | -.6000 | .0150 | 3.4383 | .1436 | .003 |

**Table 5.** False-alarm rate of the WT and Peak-to-Peak methods for the Names experiment. presents the outcomes of the false-alarm test for the innocent group in the Names experiment, using both methods. Columns (**P1-P6**) refer to the six permutations that arose from changing the role between the three identical conditions of Probe, Irrelevant1, and Irrelevant2. At an alpha level of .05, only one false alarm was obtained by the WT, compared with four by the Peak-to-Peak (PP) method. The false-positive alarms are indicated with asterisks.

| Participant | P1 |  | P2 |  | P3 |  | P4 |  | P5 |  | P6 |  |
| --- | --- | --- | --- | --- | --- | --- | --- | --- | --- | --- | --- | --- |
|  | WT | PP | WT | PP | WT | PP | WT | PP | WT | PP | WT | PP |
| 1 | .969 | .718 | .324 | .256 | .196 | .460 | .466 | .814 | .740 | .430 | .963 | .759 |
| 2 | .265 | .571 | .536 | .181 | .114 | **.012\*** | .056 | **.013\*** | .401 | .491 | .233 | .150 |
| 3 | .203 | .817 | .789 | .074 | .810 | .679 | .783 | .829 | .407 | .577 | .115 | .102 |
| 4 | .773 | .778 | .639 | .393 | .880 | .994 | .995 | .987 | .240 | .276 | .199 | .857 |
| 5 | .902 | .659 | .545 | .312 | .579 | .955 | .776 | .873 | .186 | .130 | .272 | .844 |
| 6 | .137 | .996 | .948 | .973 | .435 | .366 | **.015\*** | **.004\*** | .627 | .984 | .512 | .184 |
| 7 | .567 | .172 | .211 | .341 | .343 | .519 | .503 | .360 | .212 | **.010\*** | .422 | .674 |
| 8 | .489 | .441 | .535 | .351 | .187 | .177 | .262 | .585 | .851 | .915 | .704 | .477 |

**Table 6.** False-alarm rate for Birthdays experiment using the WT and Peak-to-Peak methods. lists the results of the false-alarm test using the WT and Peak-to-Peak methods, which were applied to the innocent group in the Birthdays experiment. At an alpha level of .05, one false alarm was obtained by the WT, compared with two by Peak-to-Peak (PP). The false-positive alarms are indicated with asterisks.

| Participant | WT | Peak-to-Peak |
| --- | --- | --- |
| 1 | .227 | .316 |
| 2 | **.001\*** | .379 |
| 3 | .525 | .124 |
| 4 | .877 | .138 |
| 5 | .755 | .731 |
| 6 | .991 | **.023\*** |
| 7 | .412 | .921 |
| 8 | .821 | .277 |
| 9 | .082 | **.020\*** |
| 10 | .440 | .559 |

A group-level analysis using paired t-tests of Probe against Irrelevant1 in the Names data sets resulted in significant P-values for WT, (*M* = .799, *SD* = .335, *t (14)* = 9.22, *p* = 2.5e-07), and Peak-to-Peak, (*M* = 8.16, *SD* = 6.49, *t* (14) = 4.86, *p* = .00025). WT and Peak-to-Peak values of Probe and Irrelevant1 in the Names data sets are shown in Table 3. For the Birthdays data sets, a paired t-test was applied to compare between Probe and Irrelevant at a group level. Using Peak-to-Peak, the outcomes of the paired t-test showed no significance difference between Probe and Irrelevant, (*M* = 1.0006, *SD* = 2.215, *t (11)* = 1.56, *p* = .146). In contrast, using the WT, a paired t-test resulted in a significant result, (*M* = .316, *SD* = .491, *t (11)* = 2.23, *p* = .047). WT and Peak-to-Peak values of Probe and Irrelevant in the Birthdays data sets are presented in Table 4. Figure 5 presents Pz ERPs for participants 1, 2, 8, and 14 in the Names experiment. The Probe waveform of participant1 has no clear positivity within the critical time window (300-1000 ms), which is, to some extent, similar to the Irrelevant1 waveform. Thus, neither method was able to differentiate between the two waveforms; the p-values of both methods were >.25. The null hypothesis distributions of both methods for participant 1 are shown in Figure 8. The true observed values of both methods were small and fell around the distribution’s centre, which resulted in a large p-value. Although the Probe condition for participant 2 only elicited a weak and narrow positivity at approximately 530 ms, which is absent in the Irrelevant1 ERP, both methods distinguished Probe from Irelevant1. However, the p-value of the WT (p=.011) is samller than the p-value of the Peak-to-peak measurement (p=.080). This finding arises because Probe and Fake follow a somewhat similar pattern in the defined time window. Examination of the participant’s ERPs reveals that, from approximately 550 ms to 1000 ms, the patterns of the Fake ERP and Probe are similar, and thus the WT can effectively detect the difference between the Probe and Irrelevant1.

**Figure 8.**
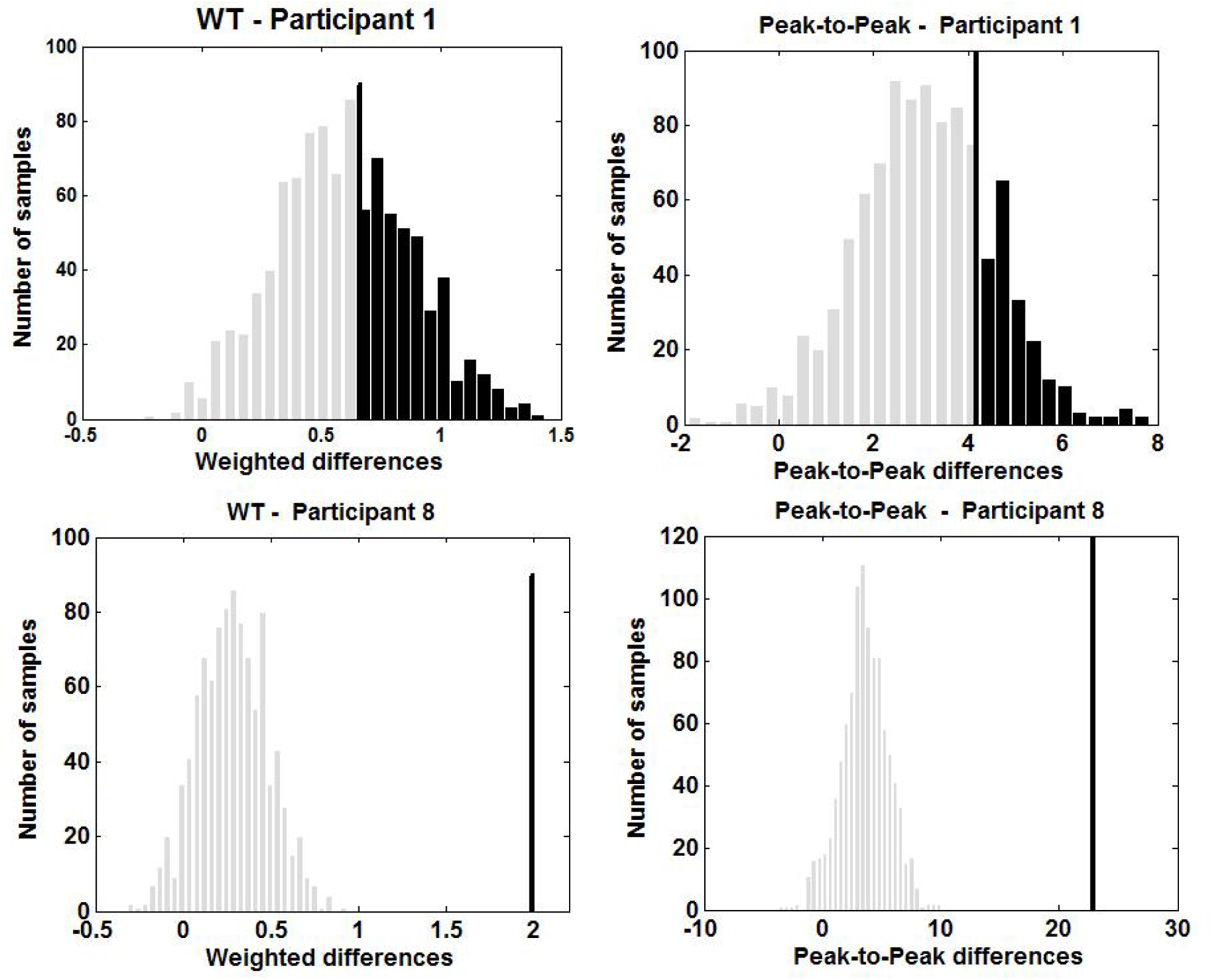
Selection of the WT and Peak-to-Peak null hypothesis distributions. Randomised null hypothesis distributions for the WT and Peak-to-Peak for two participants (1 and 8). The true observed value and Type I error region are marked in black. Participant 1, as shown in Figure 5, has a noisy Probe ERP with no evidence of P3 presence. Thus, in both methods, the true observed values were small and generated large p-values. By contrast, in both methods, participant 8, who elicited a strong P3 component, had a large true observed value, which fell outside the distribution, producing a significant p-value of <.001.

The Probe ERP of participant 8 has a clear and high positive peak (i.e., a typical P3 shape) (see Figure 5). Therefore, both of the methods detected the component well with a p-value <.001. The null hypothesis distributions for participant 8 are presented in Figure 8. While the Peak-to-Peak revealed no evidence of the presence of P3, the WT was capable of detecting participant 14 as guilty. As seen from participant 14’s ERPs in Figure 5, the Probe and Fake waveforms display atypical negative deflections at approximately 300 ms. These deflections were followed by slight respectively sharp positivity in the Probe respectively Fake waveforms. In other words, the two conditions, to some extent, display a similar pattern in the defined time window, which enables the WT to demonstrate deception for this participant; the p-values of the WT and Peak-to-Peak were .012 and .239, respectively. Figure 9 shows Pz ERPs for participants 6 and 7 from the Birthdays experiment. The Probe ERP of participant 6 has a positive deflection that peaks at approximately 550 ms post-stimulus onset (Figure 9, left). By contrast, the Irrelevant ERP has no such peak or evidence of a P3 pattern. Therefore, the WT and Peak-to-Peak measures identified the participant as guilty with p-values of .014 and .04, respectively. The Probe condition of participant 7 elicited a positive deflection, but its peak is late (around 650 ms) (Figure 9, right), which might be considered an atypical P3. Peak-to-Peak failed to identify that pattern significantly. By contrast, because the Probe and Fake ERPs are in the same positive direction between approximately 400 ms and 680 ms, the WT was able to detect the P3 response with a (significant) p-value (p=.02).

**Figure 9.**
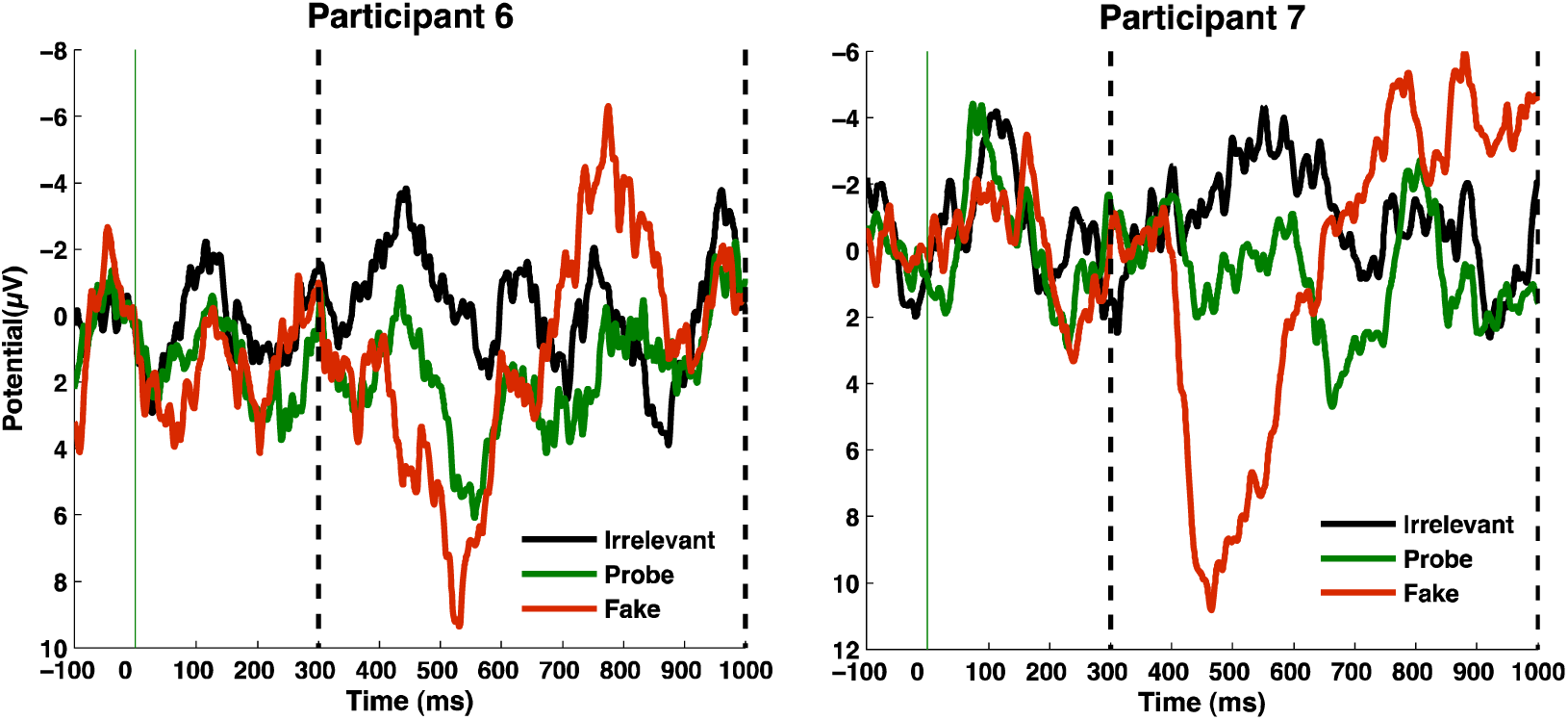
Selection of Pz ERPs from the Birthdays experiment. ERPs at Pz for participants 6 and 7 from the Birthdays experiment. Positive is down, and the vertical dashed lines mark the region in which we search for the P3.

It is worth to note that the performance of the WT was also compared with the base-to-peak (b-p) measurement, which has been used in several P3-based concealed Information studies (Rosenfeld, Ward, Frigo, *et al.*, 2015; Rosenfeld, Ward, Thai, *et al.*, 2015). The b-p was measured by finding the maximally positive 100ms interval average within the critical time window (from 300 to 1000ms pot stimulus onset).

In the Names experiment, 11 participants (11/15) in the Guilty group were detected as guilty (73% hit rate) using the b-p method. The false alarm rate was 12.5% (6/48). In the Birthdays experiment, the hit rate was 17% (2/12), and the false alarm rate was 0% (0/10).

The above results showed that the WT produced more accurate classification than the Peak-to-Peak and b-p methods. As can be noted, when applied to the Names data, the WT and Peak-to-Peak methods had better performance than the b-p measurement. In the Birthdays experiment, the b-p and Peak-to-Peak methods had comparable results. However, it has been found that the Peak-to-Peak performed better than the b-p method in classifying ERPs of P3 deception detectors (Soskins, Rosenfeld and Niendam, 2001; Rosenfeld *et al.*, 2004).

## 4 Discussion

In this paper, we have proposed a simple tool for classifying deception detection ERPs based on cross-participant templating. In this way, when applied as a group-level test, identification parameters can be seen as a further hierarchical level. Using a Monte Carlo permutation test, we have evaluated the performance of our method on a number of identity deception data sets in which we sought to identify a distinguishing ERP response for a participant’s true birthday or true name. The performance of the WT was compared with that of Peak-to-Peak, which is typically used in ERP-based lie detection studies (Abootalebi et al. 2006; Bowman et al. 2013; Bowman et al. 2014; Rosenfeld et al. 2004; Rosenfeld, Ward, Frigo, et al. 2015), and is widely considered as an accurate measurement for diagnosing lie detection ERPs (Soskins, Rosenfeld and Niendam, 2001).

However, our results indicate that when applied to the Names and Birthdays data sets, the WT outperformed Peak-to-Peak in terms of both sensitivity and specificity. The hit rate using the WT was 93% and 33% for the Names and Birthdays data sets, respectively. However, the hit rates using Peak-to-Peak were 80% and 17%, respectively. In terms of the false alarm rates, using the WT, they were 2% and 10%, compared to 8% and 20% with Peak-to-Peak for the Names and Birthdays data, respectively. With the Birthdays data, both methods performed poorly because of the low SNR ERPs that were obtained during the experiment, which can be seen clearly in the grand average ERPs (see Figure 2)^2^. Thus, the low hit rate of this experimental data is not due to the small alpha level (.05) that was applied in our analysis because we re-applied our analysis at the .1 alpha level (note, a number of workers in the lie detection domain routinely use a .1 alpha level (Rosenfeld *et al.*, 2004; Meixner and Rosenfeld, 2010b; Hu *et al.*, 2012). Although this value is still low, the hit rate of the WT slightly increased to 50% (6/12), and with Peak-to-Peak, it increased to 33%. At a group level, both methods reported significant differences between Probe and Irrelevant1 in the Names data sets. For the Birthdays data sets, only the WT showed a significant difference between Probe and Irrelevant.

In a broader sense, the WT method is particularly likely to offer benefit at lower SNRs, when the Probe P3 is weakened by the relative strength of noise, making it difficult for Peak-to-Peak to detect it. This benefit of WT will particularly be the case when the Fake P3 is strong despite a relatively weak Probe P3, as often found in the lie detection setting. In this context, the Fake is effectively an explicit target, which yields a large P3b (Bowman *et al.*, 2013). Clearly though, when the Fake and Probe elicit different ERP patterns, the WT method’s advantage will weaken.

We also used simulated data to demonstrate how the sensitivity and specificity of our methods vary at different levels of SNR. ROC curve analyses indicated that the two methods display comparable performance when applied to ERP responses with well-defined positive peaks; i.e., at high SNRs. However, Peak-to-Peak is more sensitive to noise than the WT. At all SNR levels that were applied in our simulation, the WT was more stable and provided more accurate measures.

Although not reported, we have also applied the cross-correlation difference (CD) method to the Names and Birthdays data sets. This method was first introduced by (L A Farwell and Donchin, 1991) and has been used by others to assess the ERP responses of P3-based lie detection. The CD method simply compares two cross-correlation coefficients: the cross correlation of Probe (P) and Fake (F) (R1 (P, F)) and the cross correlation of P and Irrelevant (I) (R2 (P, I)). The comporison is that if R1 is greater than R2, then the Probe would be more similar to the Fake than to the Irrelevant and, therefore, the participant would be guilty. When CD was applied to our data sets, the hit rates were 33% for the Names data and 0% for the Birthdays data sets. However, its poor performance was not surprising because CD is much too conservative a measure. Indeed, Rosenfeld and colleagues have previously reported that this method is less powerful than other methods (Rosenfeld *et al.*, 2004). They indicated that, when the Fake and Probe waveforms are out of phase, the result of the cross correlation would be low even if the Probe contained a real P3 component. Thus, in this situation, CD will fail to detect the component.

From these findings, we conclude that either Peak-to-Peak or the WT will work effectively when applied to ERPs with a high SNR. In other words, their performances will be comparable when applied to ERPs that contain sharp or high positive peaks. By contrast, the WT will be more appropriate and more accurate than the other methods when applied to ERPs with low SNRs.

Although, the WT approach has been developed and illustrated here in the context of a specific application, ERP deception detection, we do believe the functional profiler idea is broadly applicable, In particular, in the same manner that functional localiser conditions are frequently incorporated into fMRI studies, functional profiler conditions could be incorporated into ERP experiments. These would be conditions known to elicit a particular component or ERP pattern, e.g., an N400 or P3. Then individual-specific profiles from such conditions could be used as templates to investigate presence of such components in experimental conditions targeted at the hypotheses of interest.

There has been some debate in the fMRI literature on the value of functional localisers. (Friston et al. 2006) Many of the points made can also be highlighted in the context of the ERP functional profile we consider here. For example, Friston et al. (2006) argued that running a pre experimental functional localiser was problematic. However, in the experiments presented here, our Fake condition (the profiler) is integrated into the main experiment, avoiding order and time effects. These might, for example, arise if a pre-experimental profiling condition were run, which, say, exhibited worse performance than the main experiment due to perceptual learning.

## Author Note

We thank Brad Wyble, Su Li and Mick Brammer for discussions that have informed the work presented here. In particular, discussion with Su Li led us to the similarity between the Weight Template method and fMRI functional localisers.

## Footnotes

1 The simulated EEG data was generated using the function *noise* described at http://www.cs.bris.ac.uk/~rafal/phasereset/

2 Both the Birthdays and Names experiments are examples of what we call the Fringe-P3 deception detection (Bowman *et al.*, 2013, 2014) SNR and Birthdays so much lower. Although still to be definitively confirmed, we believe this is because the date stimuli were presented as two parts with a fixation cross in between and participants monitored only one of the two parts looking for, say, the Fake day. As a result the Probe date, whose salience is due to the combination of both parts, is not ‘‘seen’’.

## References

Abootalebi, V., Moradi, M. H. and Khalilzadeh, M. A. (2006) ‘A comparison of methods for ERP assessment in a P300-based GKT’, International Journal of Psychophysiology, 62(2), pp. 309–320.

Bandt, C. et al. (2009) ‘A simple classification tool for single-trial analysis of ERP components.’, Psychophysiology, 46(4), pp. 747–57. doi:10.1111/j.1469-8986.2009.00816.x.

Blair, R. C. and Karniski, W. (1993) ‘An alternative method for significance testing of waveform difference potentials.’, Psychophysiology. Wiley Online Library, 30(5), pp. 518–524. Available at: http://www.ncbi.nlm.nih.gov/pubmed/8416078.

Bowman, H. et al. (2013) ‘Subliminal Salience Search Illustrated: EEG Identity and Deception Detection on the Fringe of Awareness’, PLoS ONE. edited by M. Sigman, 8(1), p. e54258. doi:10.1371/journal.pone.0054258.

Bowman, H. et al. (2014) ‘Countering countermeasures: detecting identity lies by detecting conscious breakthrough.’, PloS one, 9(3), p. e90595. doi:10.1371/journal.pone.0090595.

Brammer, M. J. et al. (1997) ‘Generic brain activation mapping in functional magnetic resonance imaging: a nonparametric approach.’, Magnetic Resonance Imaging, 15(7), pp. 763–770. Available at: http://www.ncbi.nlm.nih.gov/pubmed/17668781.

Bruno Verschuere, Gershon Ben-Shakhar, E. M. (2011) Memory Detection: Theory and Application of the Concealed Information Test. Cambridge University Press.

Carpenter, J. and Bithell, J. (2000) ‘Bootstrap confidence intervals: when, which, what? A practical guide for medical statisticians.’, Statistics in medicine, 19(9), pp. 1141–64. doi:10.1002/(SICI)1097-0258(20000515)19:9<1141::AID-SIM479>3.0.CO;2-F-id>.

Drongelen, W. van (2006) Signal Processing for Neuroscientists: An Introduction to the Analysis of Physiological Signals. Academic Press.

Egan, J. P. (1975) Signal detection theory and ROC-analysis, Academic Press. Academic Press.

Van Erkel, A. R. and Pattynama, P. M. (1998) ‘Receiver operating characteristic (ROC) analysis: basic principles and applications in radiology.’, European Journal of Radiology, 27(2), pp. 88–94. Available at: http://www.ncbi.nlm.nih.gov/pubmed/9639133.

Farwell, L A and Donchin, E. (1991) ‘The truth will out: interrogative polygraphy (“lie detection”) with event-related brain potentials.’, Psychophysiology, 28(5), pp. 531–547. doi:10.1111/j.1469-8986.1991.tb01990.x.

Farwell, Lawrence A. and Donchin, E. (1991) ‘The Truth Will Out: Interrogative Polygraphy (“Lie Detection”) With Event-Related Brain Potentials’, Psychophysiology, 28(5), pp. 531–547. doi:10.1111/j.1469-8986.1991.tb01990.x.

Farwell, L. A., Richardson, D. C. and Hernandez, R. S. (2006) ‘Brain fingerprinting in field conditions’, in Psychophysiology, pp. S38--S38.

Filetti, M. (2013) ‘Exploring subliminal salience using the P-300□: applications to identity deception’. University of Kent.

Friston, K. J. et al. (2006) ‘A critique of functional localisers.’, NeuroImage, 30(4), pp. 1077–1087.

Groppe, D. M., Urbach, T. P. and Kutas, M. (2011) ‘Mass univariate analysis of event-related brain potentials/fields I: a critical tutorial review.’, Psychophysiology. Blackwell Publishing Inc, 48(12), pp. 1711–25. doi:10.1111/j.1469-8986.2011.01273.x.

Hu, X. et al. (2012) ‘Increasing the number of irrelevant stimuli increases ability to detect countermeasures to the P300-based Complex Trial Protocol for concealed information detection.’, Psychophysiology, 49(1), pp. 85–95. doi:10.1111/j.1469-8986.2011.01286.x.

Kilner, J. M. (2013) ‘Bias in a common EEG and MEG statistical analysis and how to avoid it.’, Clinical neurophysiology□: official journal of the International Federation of Clinical Neurophysiology, 124(10), pp. 2062–3. doi:10.1016/j.clinph.2013.03.024.

Labkovsky, E. and Rosenfeld, J Peter (2012) ‘The P300-based, complex trial protocol for concealed information detection resists any number of sequential countermeasures against up to five irrelevant stimuli.’, Applied Psychophysiology and Biofeedback, 37(1), pp. 1–10. doi:10.1007/s10484-011-9171-0.

Labkovsky, E. and Rosenfeld, J. Peter (2012) ‘The P300-based, complex trial protocol for concealed information detection resists any number of sequential countermeasures against up to five irrelevant stimuli’, Applied Psychophysiology Biofeedback, 37(1), pp. 1–10.

Meixner, J. B. et al. (2013) ‘P900: a putative novel ERP component that indexes countermeasure use in the P300-based concealed information test.’, Applied psychophysiology and biofeedback, 38(2), pp. 121–32. doi:10.1007/s10484-013-9216-7.

Meixner, J. B. and Rosenfeld, J. P. (2010a) ‘A mock terrorism application of the P300-based concealed information test.’, Psychophysiology, 48, pp. 149–154. doi:10.1111/j.1469-8986.2010.01050.x.

Meixner, J. B. and Rosenfeld, J. P. (2010b) ‘Countermeasure mechanisms in a P300-based concealed information test.’, Psychophysiology, 47(1), pp. 57–65. doi:10.1111/j.1469-8986.2009.00883.x.

Rosenfeld, J. P. et al. (2004) ‘Simple, effective countermeasures to P300-based tests of detection of concealed information’, Psychophysiology, 41(2), pp. 205–219.

Rosenfeld, J. P., Ward, A., Frigo, V., et al. (2015) ‘Evidence suggesting superiority of visual (verbal) vs. auditory test presentation modality in the P300-based, Complex Trial Protocol for concealed autobiographical memory detection’, International Journal of Psychophysiology. Elsevier, 96(1), pp. 16–22.

Rosenfeld, J. P., Ward, A., Thai, M., et al. (2015) ‘Superiority of pictorial versus verbal presentation and initial exposure in the p300-based, complex trial protocol for concealed memory detection.’, Applied psychophysiology and biofeedback. Springer US, 40(2), pp. 61–73. doi:10.1007/s10484-015-9275-z.

Soskins, M., Rosenfeld, J. P. and Niendam, T. (2001) ‘Peak-to-peak measurement of P300 recorded at 0.3 Hz high pass filter settings in intraindividual diagnosis: complex vs. simple paradigms.’, International journal of psychophysiology□: official journal of the International Organization of Psychophysiology, 40(2), pp. 173–80. Available at: http://www.ncbi.nlm.nih.gov/pubmed/11165356.

Wasserman, S. and Bockenholt, U. (1989) ‘Bootstrapping: Applications to Psychophysiology’, Psychophysiology, 26(2), pp. 208–221. doi:10.1111/j.1469-8986.1989.tb03159.x.

Yeung, N. et al. (2004) ‘Detection of synchronized oscillations in the electroencephalogram: An evaluation of methods’, Psychophysiology, 41(6), pp. 822–832. doi:10.1111/j.1469-8986.2004.00239.x.

Yeung, N. et al. (2007) ‘Theta phase resetting and the error-related negativity.’, Psychophysiology, 44(1), pp. 39–49. Available at: http://www.ncbi.nlm.nih.gov/pubmed/17241139.

