## supplemental for "Countering Cross-Individual Variance in Event Related Potentials with Functional Profiling"

### Appendix A

Table A.1 P-values by SNR.

Table A.1 presents the p-values for each method across the 15 participants in the Names experiment at different levels of SNR. The SNR values were estimated as the average of the SNRs of both conditions: Probe and Irrelevant1. Each p-value was obtained by taking the average of 100 p-values that arose from 100 iterations of generating random noise data. As *K* increases, the SNR declines, and therefore the p-value increases to reach .5. The two methods performed almost equally at a high SNR. However, at low SNRs, the p-values of the WT across most of the participants were significantly smaller than the corresponding values for Peak-to-Peak (PP). Note the K refers to the noise amplitude.

| **K** | **Participant 1** | | | **Participant 2** | | | **Participant 3** | | |
|---|---|---|---|---|---|---|---|---|---|
| | **SNR** | **WT** | **PP** | **SNR** | **WT** | **PP** | **SNR** | **WT** | **PP** |
| **1** | **.1370** | <. 0001 | <. 0001 | **.1390** | <. 0001 | <. 0001 | **.1329** | <. 0001 | <. 0001 |
| **7** | **.0400** | .0097 | .0041 | **.0360** | <. 0001 | <. 0001 | **.0379** | <. 0001 | <. 0001 |
| **13** | **.0150** | .1377 | .1357 | **.0130** | .0034 | .0242 | **.0146** | .0000 | .0001 |
| **19** | **.0070** | .2361 | .2416 | **.0060** | .0268 | .1267 | **.0074** | .0022 | .0039 |
| **25** | **.0040** | .3858 | .4052 | **.0040** | .0752 | .2188 | **.0044** | .0202 | .0326 |
| **31** | **.0030** | .4020 | .4222 | **.0025** | .1469 | .3171 | **.0029** | .0440 | .0672 |
| **37** | **.0020** | .3710 | .3977 | **.0017** | .1979 | .3630 | **.0021** | .1024 | .1579 |
| **43** | **.0015** | .4861 | .4980 | **.0015** | .2043 | .3328 | **.0016** | .1049 | .1603 |
| **49** | **.0010** | .4185 | .4525 | **.0010** | .2503 | .3825 | **.0012** | .1715 | .2160 |
| **K** | **Participant 4** | | | **Participant 5** | | | **Participant 6** | | |
| | **SNR** | **WT** | **PP** | **SNR** | **WT** | **PP** | **SNR** | **WT** | **PP** |
| **1** | **.1352** | <.0001 | <.0001 | **.1321** | <.0001 | <.0001 | **.1285** | <.0001 | <.0001 |
| **7** | **.0352** | <.0001 | <.0001 | **.0411** | <.0001 | <.0001 | **.0585** | <.0001 | <.0001 |
| **13** | **.0125** | .0004 | .0001 | **.0166** | .0003 | <.0001 | **.0329** | <.0001 | <.0001 |
| **19** | **.0062** | .0104 | .0184 | **.0085** | .0008 | .0005 | **.0198** | <.0001 | <.0001 |
| **25** | **.0036** | .0480 | .0621 | **.0052** | .0116 | .0131 | **.0127** | .0035 | <.0001 |
| **31** | **.0024** | .0877 | .1161 | **.0034** | .0370 | .0447 | **.0088** | .0020 | .0001 |
| **37** | **.0017** | .1232 | .1552 | **.0024** | .0589 | .0566 | **.0064** | .0110 | .0018 |
| **43** | **.0013** | .1667 | .2057 | **.0018** | .0993 | .1249 | **.0048** | .0093 | .0077 |
| **49** | **.0010** | .2056 | .2881 | **.0014** | .1093 | .1688 | **.0038** | .0236 | .0200 |
| **K** | **Participant 7** | | | **Participant 8** | | | **Participant 9** | | |
| | **SNR** | **WT** | **PP** | **SNR** | **WT** | **PP** | **SNR** | **WT** | **PP** |
| **1** | **.1296** | <.0001 | <.0001 | **.1384** | <.0001 | <.0001 | **.1223** | <.0001 | <.0001 |
| **7** | **.0359** | <.0001 | <.0001 | **.0589** | <.0001 | <.0001 | **.0274** | <.0001 | <.0001 |
| **13** | **.0138** | <.0001 | <.0001 | **.0278** | <.0001 | <.0001 | **.0099** | <.0001 | <.0001 |
| **19** | **.0070** | .0016 | .0022 | **.0153** | <.0001 | <.0001 | **.0050** | .0045 | .0105 |
| **25** | **.0041** | .0243 | .0329 | **.0095** | .0005 | <.0001 | **.0029** | .0283 | .0615 |
| **31** | **.0027** | .0349 | .0614 | **.0064** | .0007 | .0003 | **.0019** | .0681 | .1198 |
| **37** | **.0019** | .0634 | .0953 | **.0046** | .0044 | .0060 | **.0014** | .0806 | .1636 |
| **43** | **.0015** | .0594 | .1012 | **.0035** | .0069 | .0085 | **.0010** | .1266 | .2066 |
| **49** | **.0011** | .1158 | .1908 | **.0027** | .0137 | .0231 | **.0008** | .1608 | .2976 |

| K | Participant 10 | | | Participant 11 | | | Participant 12 | | |
|---|---|---|---|---|---|---|---|---|---|
| | SNR | WT | PP | SNR | WT | PP | SNR | WT | PP |
| **1** | **.1525** | <.001 | <.001 | **.1335** | <.001 | <.001 | **.1446** | <.001 | <.001 |
| **7** | **.0433** | <.001 | <.001 | **.0333** | <.001 | <.001 | **.0639** | <.001 | <.001 |
| **13** | **.0159** | .0010 | .0016 | **.0120** | .0012 | .0148 | **.0280** | <.001 | <.001 |
| **19** | **.0079** | .0299 | .0381 | **.0060** | .0239 | .0823 | **.0150** | .0007 | .0002 |
| **25** | **.0046** | .0667 | .1045 | **.0035** | .0778 | .1642 | **.0091** | .0089 | .0054 |
| **31** | **.0031** | .1304 | .1735 | **.0023** | .1378 | .2236 | **.0061** | .0278 | .0286 |
| **37** | **.0022** | .1919 | .2507 | **.0017** | .1852 | .2844 | **.0044** | .0714 | .0698 |
| **43** | **.0016** | .2183 | .2550 | **.0012** | .2115 | .2976 | **.0033** | .0680 | .0673 |
| **49** | **.0012** | .2867 | .3278 | **.0009** | .3070 | .3886 | **.0026** | .1177 | .1437 |

| K | Participant 13 | | | Participant 14 | | | Participant 15 | | |
|---|---|---|---|---|---|---|---|---|---|
| | SNR | WT | PP | SNR | WT | PP | SNR | WT | PP |
| **1** | **.1416** | <.001 | <.001 | **.1284** | <.001 | <.001 | **.1438** | <.001 | <.001 |
| **7** | **.0391** | <.001 | <.001 | **.0245** | <.001 | .0051 | **.0299** | <.001 | <.001 |
| **13** | **.0148** | .0002 | .0001 | **.0082** | .0030 | .0955 | **.0106** | .0015 | .0010 |
| **19** | **.0073** | .0101 | .0041 | **.0039** | .0309 | .1723 | **.0053** | .0382 | .0542 |
| **25** | **.0043** | .0286 | .0345 | **.0023** | .1081 | .3329 | **.0031** | .0741 | .1200 |
| **31** | **.0029** | .0644 | .0635 | **.0015** | .1825 | .3749 | **.0021** | .1486 | .1888 |
| **37** | **.0020** | .1015 | .1061 | **.0011** | .2265 | .4016 | **.0014** | .1621 | .2313 |
| **43** | **.0015** | .1866 | .2162 | **.0008** | .2760 | .4247 | **.0011** | .2066 | .2537 |
| **49** | **.0012** | .1947 | .1782 | **.0006** | .3216 | .4651 | **.0008** | .1900 | .2530 |

Table A.2 AUCs by SNR.

Table A.2 list the areas under the ROC curve for each method across all of the participants in the Names experiment at different levels of SNR. As K increases, the SNR decreases, and therefore the AUC decreases. The two methods performed almost equally well at a high SNR. However, at low SNRs, the AUCs of the WT were consistently higher than the corresponding AUCs of the Peak-to-Peak (PP). Note that K refers to the noise amplitude.

| K | Participant 1 | | | Participant 2 | | | Participant 3 | | |
|---|---|---|---|---|---|---|---|---|---|
| | **SNR** | **WT** | **PP** | **SNR** | **WT** | **PP** | **SNR** | **WT** | **PP** |
| **1** | **.1370** | 1.0000 | 1.0000 | **.1390** | 1.000 | 1.0000 | **.1329** | 1.0000 | 1.000 |
| **7** | **.0400** | .9872 | .9972 | **.0360** | 1.000 | 1.0000 | **.0379** | 1.0000 | 1.000 |
| **13** | **.0150** | .8493 | .8697 | **.0130** | .9946 | .9774 | **.0146** | .9999 | 1.000 |
| **19** | **.0070** | .7439 | .7589 | **.0060** | .9708 | .8770 | **.0074** | .9960 | .9980 |
| **25** | **.0040** | .5782 | .5933 | **.0040** | .9174 | .7833 | **.0044** | .9770 | .9686 |
| **31** | **.0030** | .5621 | .5782 | **.0025** | .8384 | .6841 | **.0029** | .9527 | .9355 |
| **37** | **.0020** | .5960 | .6019 | **.0017** | .7816 | .6363 | **.0021** | .8862 | .8438 |
| **43** | **.0015** | .4728 | .4977 | **.0015** | .7766 | .6664 | **.0016** | .8850 | .8402 |
| **49** | **.0010** | .5454 | .5441 | **.0010** | .7247 | .6173 | **.0012** | .8113 | .7842 |
| **K** | **Participant 4** | | | **Participant 5** | | | **Participant 6** | | |
| | **SNR** | **WT** | **PP** | **SNR** | **WT** | **PP** | **SNR** | **WT** | **PP** |
| **1** | **.1352** | 1.0000 | 1.0000 | **.1321** | 1.0000 | 1.0000 | **.1285** | 1.000 | 1.000 |
| **7** | **.0352** | 1.0000 | 1.0000 | **.0411** | 1.0000 | 1.0000 | **.0585** | 1.000 | 1.000 |
| **13** | **.0125** | .9992 | 1.0000 | **.0166** | .9998 | 1.0000 | **.0329** | 1.000 | 1.000 |
| **19** | **.0062** | .9875 | .9841 | **.0085** | .9983 | .9999 | **.0198** | 1.000 | 1.000 |
| **25** | **.0036** | .9464 | .9410 | **.0052** | .9857 | .9888 | **.0127** | .9954 | 1.000 |
| **31** | **.0024** | .9025 | .8868 | **.0034** | .9587 | .9583 | **.0088** | .9972 | 1.000 |
| **37** | **.0017** | .8646 | .8467 | **.0024** | .9356 | .9465 | **.0064** | .9869 | .9991 |
| **43** | **.0013** | .8179 | .7953 | **.0018** | .8909 | .8775 | **.0048** | .9882 | .9934 |
| **49** | **.0010** | .7733 | .7127 | **.0014** | .8814 | .8326 | **.0038** | .9732 | .9814 |
| **K** | **Participant 7** | | | **Participant 8** | | | **Participant 9** | | |
| | **SNR** | **WT** | **PP** | **SNR** | **WT** | **PP** | **SNR** | **WT** | **PP** |
| **1** | **.1296** | 1.0000 | 1.0000 | **.1384** | 1.0000 | 1.0000 | **.1223** | 1.000 | 1.000 |
| **7** | **.0359** | 1.0000 | 1.0000 | **.0589** | 1.0000 | 1.0000 | **.0274** | 1.000 | 1.000 |
| **13** | **.0138** | .9999 | 1.0000 | **.0278** | 1.0000 | 1.0000 | **.0099** | 1.000 | 1.000 |
| **19** | **.0070** | .9978 | .9985 | **.0153** | 1.0000 | 1.0000 | **.0050** | .9938 | .9899 |
| **25** | **.0041** | .9718 | .9700 | **.0095** | .9999 | 1.0000 | **.0029** | .9674 | .9393 |
| **31** | **.0027** | .9607 | .9419 | **.0064** | .9986 | 1.0000 | **.0019** | .9245 | .8805 |
| **37** | **.0019** | .9303 | .9080 | **.0046** | .9946 | .9953 | **.0014** | .9105 | .8349 |
| **43** | **.0015** | .9360 | .9012 | **.0035** | .9909 | .9922 | **.0010** | .8604 | .7954 |
| **49** | **.0019** | .8721 | .8097 | **.0046** | .9841 | .9800 | **.0008** | .8220 | .6992 |

| K | Participant 10 | | | Participant 11 | | | Participant 12 | | |
|---|---|---|---|---|---|---|---|---|---|
| | SNR | WT | PP | SNR | WT | PP | SNR | WT | PP |
| **1** | **.1525** | 1.000 | 1.000 | **.1335** | 1.000 | 1.000 | **.1446** | 1.000 | 1.000 |
| **7** | **.0433** | 1.000 | 1.000 | **.0333** | 1.000 | 1.000 | **.0639** | 1.000 | 1.000 |
| **13** | **.0159** | .9979 | .9994 | **.0120** | .9975 | .9873 | **.0280** | 1.000 | 1.000 |
| **19** | **.0079** | .9660 | .9625 | **.0060** | .9722 | .9195 | **.0150** | .9985 | 1.000 |
| **25** | **.0046** | .9274 | .8984 | **.0035** | .9133 | .8362 | **.0091** | .9892 | .9956 |
| **31** | **.0031** | .8569 | .8300 | **.0023** | .8481 | .7773 | **.0061** | .9687 | .9725 |
| **37** | **.0022** | .7892 | .7511 | **.0017** | .7961 | .7172 | **.0044** | .9216 | .9335 |
| **43** | **.0016** | .7597 | .7483 | **.0012** | .7676 | .7032 | **.0033** | .9251 | .9353 |
| **49** | **.0012** | .6855 | .6715 | **.0009** | .6664 | .6085 | **.0026** | .8709 | .8568 |

| K | Participant 13 | | | Participant 14 | | | Participant 15 | | |
|---|---|---|---|---|---|---|---|---|---|
| | SNR | WT | PP | SNR | WT | PP | SNR | WT | PP |
| **1** | **.1416** | 1.000 | 1.000 | **.1284** | 1.000 | 1.000 | **.1438** | 1.000 | 1.000 |
| **7** | **.0391** | 1.000 | 1.000 | **.0245** | 1.000 | .9960 | **.0299** | 1.000 | 1.000 |
| **13** | **.0148** | .9992 | 1.000 | **.0082** | .9945 | .9078 | **.0106** | .9970 | .9998 |
| **19** | **.0073** | .9871 | .9970 | **.0039** | .9656 | .8295 | **.0053** | .9574 | .9484 |
| **25** | **.0043** | .9672 | .9679 | **.0023** | .8819 | .6670 | **.0031** | .9181 | .8822 |
| **31** | **.0029** | .9293 | .9376 | **.0015** | .7991 | .6251 | **.0021** | .8379 | .8120 |
| **37** | **.0020** | .8891 | .8962 | **.0011** | .7536 | .5944 | **.0014** | .8225 | .7674 |
| **43** | **.0015** | .7959 | .7851 | **.0008** | .6995 | .5748 | **.0011** | .7727 | .7477 |
| **49** | **.0012** | .7865 | .8244 | **.0006** | .6445 | .5325 | **.0008** | .7919 | .7470 |

#### Appendix B

##### The staircase procedure

Since the optimal SOA for date stimuli shown in our RSVP format depends on the participant, a staircase test of perception threshold was employed. The optimal SOA was defined as the lowest SOA for which the participant stayed under the set error threshold (specified below). The test comprised 2 types of RSVP trials: *Target* and *Irrelevant.* Prior to the start of the test, participants were given a target date, which only appeared in *Target* trials. The position of the target in the stream was selected pseudorandomly: from 5 to 10. Every RSVP stream was followed by a question (shown after the *last* item question) that asked the participant whether they saw the target or not; their accuracy in responding to this question appropriately was recorded. The structure of the test can be summarised by the following algorithm:

1) **Run 7 trials (3 Target trials and 4 Irrelevant trials, randomly mixed) with an SOA of 400ms.**
2) **Repeatedly decrease the SOA by 100ms until hit rate < 74%, or false alarm rate > 21%.**
3) **Repeatedly increase the SOA by 50ms until hit rate ≥ 75%, or false alarm rate ≤ 20%.**
4) **Run 14 trials (6 Target trials and 8 Irrelevant trials, randomly mixed) with the final SOA from step 3.**
5) **If hit rate > 84 % or false alarm rate < 12 % repeat step 4 with SOA - 25 ms.**
6) **If hit rate ≤ 68 % or false alarm rate ≥ 24 %, repeat step 4 with SOA + 25 ms. If not, return current SOA as optimal**
